# Modulation of RNA-dependent interactions in stress granules prevents persistent TDP-43 accumulation in ALS/FTD

**DOI:** 10.1101/474577

**Authors:** Mark Y. Fang, Sebastian Markmiller, William E. Dowdle, Anthony Q. Vu, Paul J. Bushway, Sheng Ding, Mark M. Mercola, Joseph W. Lewcock, Gene W. Yeo

## Abstract

Human genetic variants are usually represented by four values with variable length: chromosome, position, reference and alternate alleles. Thereis no guarantee that these components are represented in a consistent way across different data sources, and processing variant-based data can be inefficient because four different comparison operations are needed for each variant, three of which are string comparisons. Working with strings, in contrast to numbers, poses extra challenges on computer memory allocation and data-representation. Existing variant identifiers do not typicallyrepresent every possible variant we may be interested in, nor they are directly reversible. To overcome these limitations, *VariantKey*, a novel reversible numerical encoding schema for human genetic variants, is presented here alongside a multi-language open-source software implementation (http://github.com/genomicspls/variantkey). VariantKey represents variants as single 64 bit numeric entities, while preserving the ability to be searched and sorted by chromosome and position. The individual components of short variants can be directly read back from the VariantKey, while long variants are supported with a fast lookup table.

**Highlights:**

1. ~100 compounds identified by high-content screen inhibit SGs in HEK293, NPCs and iPS-MNs.
2. ALS-associated RBPs are recruited to SGs in an RNA-dependent manner
3. Molecules with planar moieties prevent recruitment of ALS-associated RBPs to SGs
4. Compounds inhibit TDP-43 accumulation in SGs and in *TARDBP* mutant iPS-MNs.

## INTRODUCTION

Stress granules (SGs) assemble transiently in response to cellular stresses such as oxidative damage, heat shock and environmental toxins, as an adaptive survival mechanism for cells (*1, 2*). SGs contain both proteins as well as cellular messenger RNAs, which are locked in a translationally stalled state induced by phosphorylation of serine 51 of the translation initiation factor eIF2α (*3, 4*). By modulating mRNA translation and recruiting intracellular signaling proteins, SGs are believed to triage intracellular activity toward an integrated stress response (*5–8*). SGs are highly dynamic, exhibiting liquid-like behaviors and begin to disassemble within minutes of removal of stress (*9*). These liquid-like properties are thought to be mediated by the intrinsically disordered regions (IDRs) common to many proteins found in SGs (*10–15*). Neurodegeneration-linked mutations in genes such as *FUS*, *HNRNPA2B1 and TARDBP* (encoding TDP-43), frequently cluster in areas corresponding to IDRs in the encoded proteins, potentially altering their liquid-like phase-separation properties (*16–18*). These mutations are implicated in hereditary forms of frontotemporal dementia (FTD) and amyotrophic lateral sclerosis (ALS) (*19–23*), a fatal, incurable disease characterized by progressive degeneration of motor neurons (MNs) (*24*). *In vitro* studies of recombinant IDRs carrying ALS-associated mutations report that phase-separated droplets of mutant IDRs can transition from a liquid-like state to a solid-like state resembling prion inclusion bodies and characterized by cross-beta sheet fibrils (*18, 19, 25-27*). To illustrate, recombinant mutant IDR from HNRNPA2B1 undergoes LLPS followed by spontaneous maturation into insoluble fibers (*18, 19*). Therefore, these IDR mutations likely predispose assembly of inclusion bodies, and is speculated to cause toxic loss-of-function and contributing to disease pathophysiology. Indeed, a hallmark feature of nearly all ALS patients is the presence of cytoplasmic TDP-43-containing inclusion bodies within MNs that contain SG-associated proteins such as PABPC1, TIA1, eIF3, ATXN2, HNRNPA2B1, and FUS (*19, 28–32*).

Recent studies that characterize the composition of SGs have revealed that a large fraction of protein components found in SG appear to be pre-assembled prior to stress (*11, 12*). Also, a super-resolution microscopy study reported the existence of substructures called SG cores, around which a halo of additional proteins forms, called the SG shell (*10*). It is very likely that cores and shells contain different protein components, with differences that may relate to pathogenesis of disease (*3, 10*). Excitingly, the modulation of some SG protein components appears to alleviate degenerative phenotypes in animal models of ALS (*11, 33–35*). Despite these advances, there still exists an urgent need to understand how ALS-associated proteins such as TDP-43 relate to SGs and for new tools that can readily perturb these relationships.

Thus, to accelerate our understanding of SGs and their connections to molecular pathophysiology in neurodegenerative disease, we conducted a high-content screen for small molecules that robustly modulate various aspects of SG biology. We identified several classes of compounds, including small molecules that act at cell surface targets such as ion channels, receptors, or lipid membranes, and compounds that modulate inflammatory signaling pathways. Interestingly, we also identified a large group of molecules with planar moieties such as nucleic acid intercalators, supporting the hypothesis that SG formation requires nucleic acid interactions. In parallel, we discovered that SGs accumulate ALS-associated proteins in an RNA-dependent manner, and that molecules with planar moieties disrupt this accumulation. We show SGs can induce persistent cytoplasmic localization of TDP-43 in an induced pluripotent stem cell-derived MN (iPS-MN) model of disease, and that iPS-MNs carrying ALS-associated mutations in *TARDBP* or FUS exhibit exacerbated cytoplasmic localization of TDP-43. Excitingly, nucleic acid intercalators with planar moieties prevent co-localization of TDP-43 with SGs and reduce cytoplasmic localization of TDP-43 in mutant iPS-MNs. We believe our results expand our understanding of SG biology, establish a mechanistic relationship from SGs to disease pathophysiology, and pave the way to development of a new classes of therapeutics for ALS/FTD.

## RESULTS

### Generation of Robust SG Assays

G3BP1 is a key protein required for SG formation (*1, 36*). As G3BP1 is found in both the SG core and, to a lesser extent, the SG shell (*10*), we reasoned that identifying compounds that modulate G3BP1-positive puncta formation will unveil molecular principles of SG formation and point toward mechanistic links between SGs and disease pathophysiology. We used CRISPR/Cas9 genome editing in HEK293xT cells and the human induced pluripotent stem cell iPSC line CV-B to insert the coding sequence of green fluorescent protein (GFP) immediately upstream of the stop codon of the endogenous *G3BP1* locus. iPSCs were subsequently differentiated into neural precursor cells using a small molecule protocol (smNPCs) or motor neurons (MNs; Figures 1A-B and S1A). The G3BP1-GFP expressing HEK293xT cells, NPCs, and iPS-MNs robustly and reproducibly formed bright GFP-positive(+) puncta upon exposure to sodium arsenite (NaAsO_2_), consistent with formation of G3BP1-containing SGs (Figure 1C). These NaAsO2-induced puncta were abrogated by pre-treatment with cycloheximide, which inhibits SG formation by stabilizing polysomes, as expected (Figures 1C-D) (*1*). Our results indicate that these G3BP1-GFP expressing lines are robust SG reporter lines.

**Figure 1.**
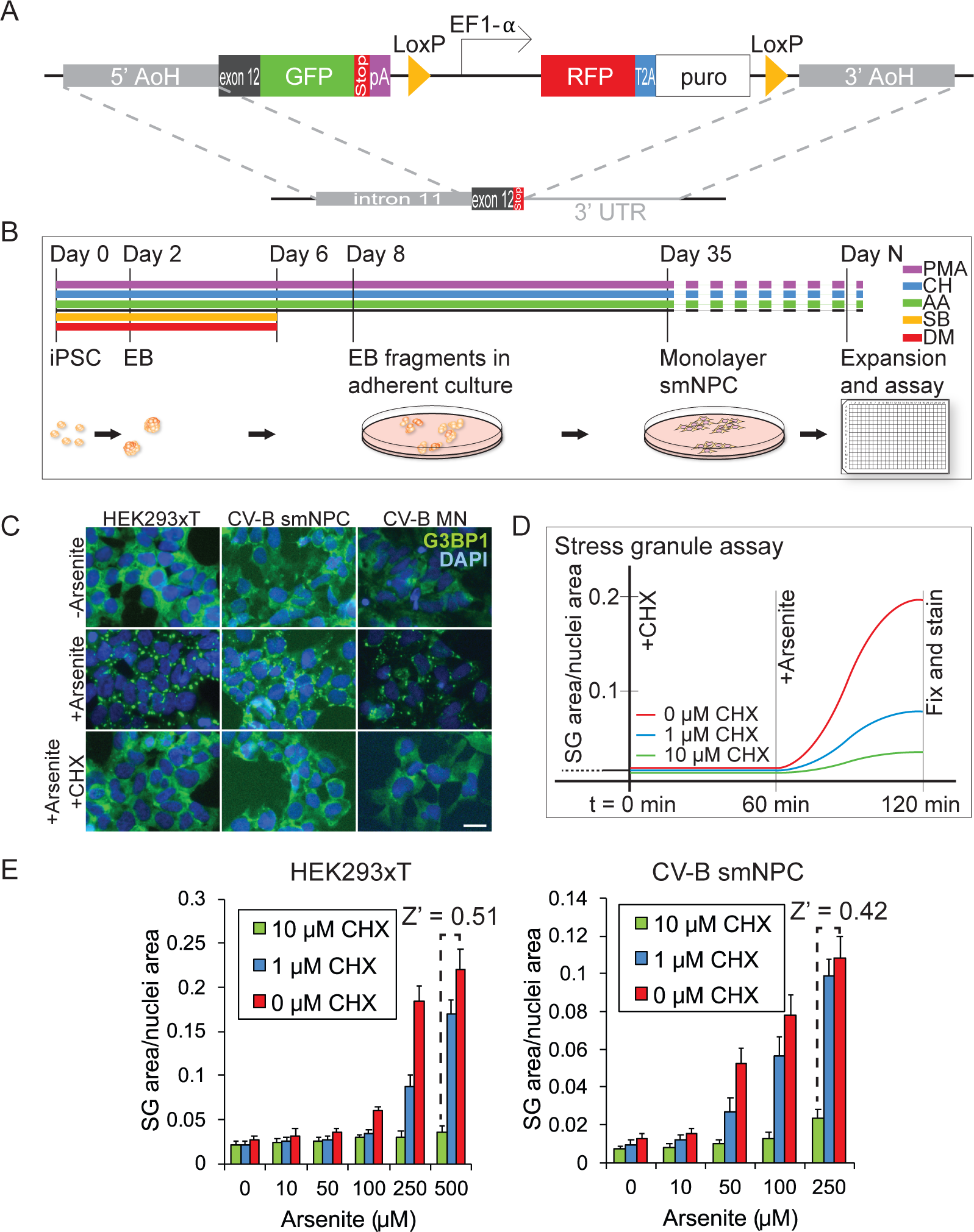
Generation of SG reporter lines. (A) Schematic showing the GFP construct inserted in the C-terminus of the endogenous G3BP1 gene locus using CRISPR/Cas9-mediated homology directed repair. The puromycin selection cassette was removed by Cre-mediated recombination of flanking LoxP sequences. AoH: arm of homology, pA: polyadenylation sequence, puro: puromycin resistance cassette, UTR: untranslated region. (B) Timeline for the generation of small molecule neural precursor cells used in screening. iPSC: induced-pluripotent stem cells, EB: embryoid bodies, NPC: small molecule neural precursor cells, PMA: Purmorphamine, CH: CHIR99021, AA: L-Ascorbic acid, SB: SB431542, DM: Dorsomorphin. (C) Representative wide-field fluorescence microscopy of G3BP1-GFP SG reporter lines under unstressed, NaAsO_2_ -stressed, and stressed but with cycloheximide pretreatment conditions. Scale bar is 50 μm. MN: motor neurons. (D) Overview of the screening paradigm, in which cells are pretreated with compounds such as cycloheximide for one hour, after which NaAsO_2_ is added to stress cells for one hour. CHX: cycloheximide. (E) Histograms showing SG area/nuclei area at varying NaAsO_2_ concentrations in the absence and presence of cycloheximide pretreatment, a positive control SG inhibiting compound. The Z-factors (Z’) for the screen assay conditions are shown in the upper right and are used to estimate whether the screen assays have adequate sensitivity and specificity for high-throughput screening: 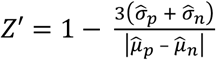, where 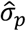 and 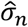 are the sample standard deviations for the positive and negative controls, respectively, and 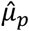 and 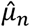 are the sample means for the positive and negative controls, respectively. Error bars are sample standard deviations from four biological replicates.

We conducted our primary SG screen in the HEK293xT and NPC reporter lines, as they can be expanded to large quantities and seeded in a monodisperse layer facilitating high-content image segmentation. The screening strategy entailed first pre-incubating cells with compounds before stressing the cells with NaAsO_2_ (Figure 1D). We performed the screen with a compound-first, stress-second paradigm because this order of operations is most likely to capture both compounds that prevent SGs from forming as well as compounds that cause disassembly of SGs while they have already begun to assemble. Using cycloheximide as a positive control and DMSO vehicle as the negative control, we calculated Z-factors of 0.51 and 0.42 for the assay in HEK293xT cells and NPCs, respectively (Figure 1E). We concluded from these Z-factors that the signal-to-noise ratio of the SG assay was sufficient for HCS (*37*).

### Identification of Diverse Classes of SG-modulating Compounds

We developed an experimental and computational pipeline to screen 3,350 and 5,910 compounds in biological duplicate in HEK293xT cells and NPCs, respectively (Figure 2A). The screened compounds were sourced from five complex small molecule libraries spanning a diverse range of chemical structures and biological activities. Cells were pre-treated with screen compounds at 10μM, then stressed with NaAsO_2_, fixed, stained with DAPI, and imaged. Screen images were segmented to identify DAPI-positive(+) nuclei and G3BP1-GFP-positive(+) SGs. To measure SG formation, the amount of SG formation per cell was quantified as the total image area enclosed in SGs divided by the total image area enclosed in nuclei. In addition to modulating the amount of SG formation per cell, these compounds may also alter the average SG size and/or the number of SGs per cell. We opined that a compound that inhibits the condensation of SG shell proteins onto SG cores may impair SG fusion and lead to on average smaller but more numerous SGs. We therefore computed two other metrics. The number of SGs formed per cell was quantified as the number of SG puncta divided by the total image area enclosed in nuclei, and the average size of SGs was quantified as the total image area enclosed in SGs divided by the number of SG puncta. After normalizing the data, we performed a modified one sample Student’s *t*-test to identify hit compounds that substantially alter one or more of these SG formation metrics.

**Figure 2.**
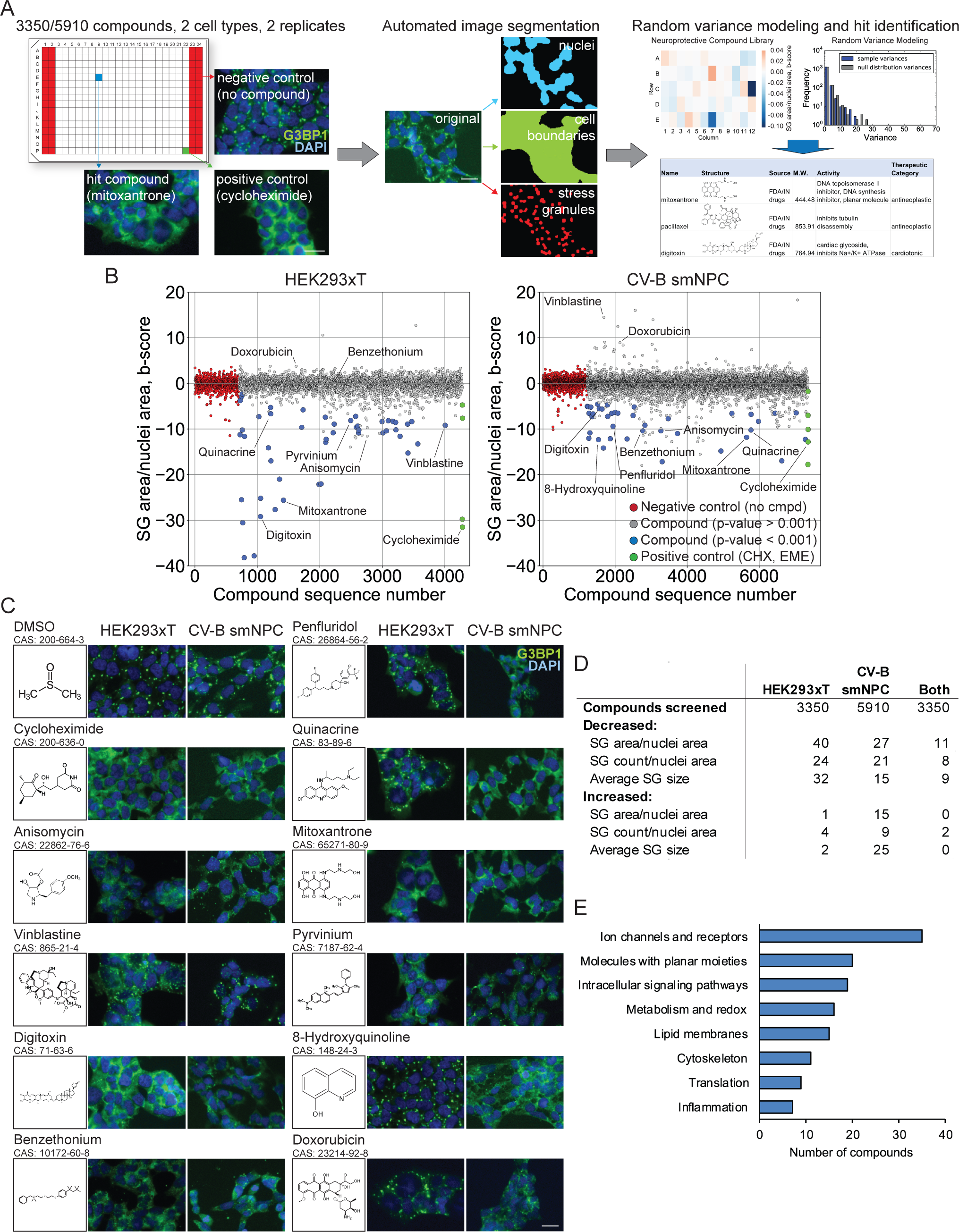
Identification of SG-modulating compounds by high-content screening. (A) Schematic depicting the experimental and computational workflow of the small molecule screen for SG-modulating compounds. Scale bar is 50 μm. (B) Scatterplot showing the amount of SG formation for each compound screened, computed as a b-score: SG area/nuclei area is processed through a two-way row/column median polish and the residuals are divided by the median absolute deviation of all post-polish residuals of the screen. For each compound, the mean of biological duplicate b-scores is represented. The red points on the left are negative controls in which no compounds were added. The green points on the right are positive controls in which either cycloheximide or emetine was added. The blue points are compounds identified as significantly reducing the amount of SG formation (p < 0.001) and the grey points are compounds that did not significantly reduce the amount of SG formation (p > 0.001). Significance levels are from one sample Student’s t-tests against a null hypothesis mean of zero. SG: stress granule, NPC: small molecule neural precursor cells, CHX: cycloheximide, EME: emetine. (C) Representative wide-field fluorescence microscopy of HEK293xT and CV-B small molecule neural precursor cell SG reporter lines treated with SG-modulating compounds. Included are the skeletal formulae of the compounds. Scale bar is 50 μm. (D) Table summarizing the number of compounds which either decreased or increased the amount of SGs per cell defined as SG area/nuclei area, the number of SGs per cell defined as SG count/nuclei area, or the average size of SGs defined as SG area/SG count. Some compounds were included in more than one library screened; these were counted only once each. (E) Histogram classifying hit compounds from the table in (D) by the known cellular targets of the compounds. Classification was performed by manual annotation from the National Center for Biotechnology Information PubChem database.

We identified 40 and 27 compounds that reduced the amount of SG formation per cell in HEK293xT cells and NPCs, respectively (Figures 2B-D and S2A). Additionally, we identified around 50 compounds which modulated SG formation in other ways, such as increasing or decreasing the average size of SGs or changing the number of SGs per cell (Figures 2D and S2A-B). Hit compounds were then annotated with the compound name, skeletal formulae, and reported cellular targets using the National Center for Biotechnology Information PubChem database (Figures 2E, S2A, and Table 1). Supporting the validity of our screening approach, we confirmed that ribosomal inhibitors, such as anisomycin, the aminoglycoside neomycin, and the macrolide roxithromycin, strongly reduced the amount of SG formation per cell (Figures 2C, 2E, S2A, and Tables 2–4). This is consistent with the fact that ribosomal inhibitors, including cycloheximide and emetine which we included in the screen as positive controls, have been previously described to affect SG formation (*1*). We also confirmed that cytoskeleton targeting compounds, such as the vinka alkaloids vincristine and vinblastine, the microtubule stabilizing drug paclitaxel, and the actin filament stabilizer prieurianin, altered average SG size and/or the number of SGs per cell, consistent with previous reports that the cytoskeleton and cytoskeleton-associated molecular motors play key roles in SG migration, fusion, and maturation (Figures 2C, 2E, S2A, and Tables 2–4) (*38*).

**Table 1.** Cellular targets of SG-modulating compounds. Table enumerating the known cellular targets of SG-modulating compounds identified in the SG screen, corresponding to the table in (Figure 2D) and the pie charts of (Figure S2A). Related cellular targets have been grouped together and cellular target superclasses have been color-coded.

|  |  |
| --- | --- |
|  | **MOLECULES WITH PLANAR MOIETIES, INTERACT WITH NUCLEIC ACIDS** |
|  | DNA methyltransferase |
|  | DNA topoisomerase |
|  | molcules with planar moieties |
|  | PARylation |
|  | **INTERACT WITH PROTEIN INTRINSICALLY DISORDERED REGIONS** |
|  | LCD/PrLD |
|  | **ION CHANNELS, RECEPTORS, TRANSPORTERS** |
|  | adrenergic receptors (alpha1, alpha2) |
|  | calcitriol receptor (vitamin D receptor) |
|  | cannabinoid receptor (CB1) |
|  | cholinergic receptors (muscarinic, nicotinic) |
|  | dopamine receptors (D2, D3, D4) |
|  | estrogen receptor |
|  | G-protein-coupled estrogen receptor 1 (GPER) |
|  | GABA-A receptor |
|  | imidazoline receptors (I1, I2) |
|  | Na<sup>+</sup>/Cl<sup>-</sup> cotransporter |
|  | Na<sup>+</sup>/K<sup>+</sup> ATPase |
|  | OATP (OATP1B1 and OATP1B3) |
|  | receptor tyrosine kinase |
|  | SERCA |
|  | serotonin receptors (5HT1A, 5HT2) |
|  | serotonin reuptake transporter (SERT) |
|  | transferrin receptor |
|  | TRPV1 |
|  | voltage gated Na<sup>+</sup> channels |
|  | **INTRACELLULAR SIGNALING, TRANSCRIPTION FACTORS** |
|  | adenylate cyclase |
|  | Akt |
|  | CK1alpha, Wnt |
|  | DYRK |
|  | FKBP12 |
|  | HIF-1 (hypoxia inducible factor-1) |
|  | JAK-STAT |
|  | JNK/SAPK, p38 MAPK, RACK1, MTK1 |
|  | NFkappaB |
|  | PAK (p21 activated kinase) |
|  | PGC1alpha |
|  | PKC |
|  | sirtuin1 |
|  | Zn<sup>2+</sup> finger proteins |
|  | **LIPID MEMBRANES** |
|  | plasma membrane |
|  | **INFLAMMATION** |
|  | COX1, COX2 |
|  | histamine N-methyltransferase |
|  | histamine receptor (H1) |
|  | myeloperoxidase |
|  | phospholipase A2 |
|  | **METABOLISM, RESPIRATION, REDOX** |
|  | aldehyde dehydrogenase |
|  | autophagosome |
|  | carbonic anhydrase |
|  | dihydropteroate synthase |
|  | dopamine beta hydroxylase |
|  | electron transport chain |
|  | glutathione reductase and GSH |
|  | glycolytic enzymes |
|  | Hsp60/chaperonin, Hsp90 |
|  | monoamine oxidase |
|  | proteasome-chemotrypsin-like subunit |
|  | pyruvate dehydrogenase |
|  | reactive with thiols and amines |
|  | SOD2 (Mn<sup>2+</sup> dependent SOD) |
|  | **CYTOSKELETON** |
|  | microfilaments |
|  | microtubules |
|  | **TRANSLATION** |
|  | PERK |
|  | ribosome |
|  | **UNKNOWN TARGET** |

**Table 2.**
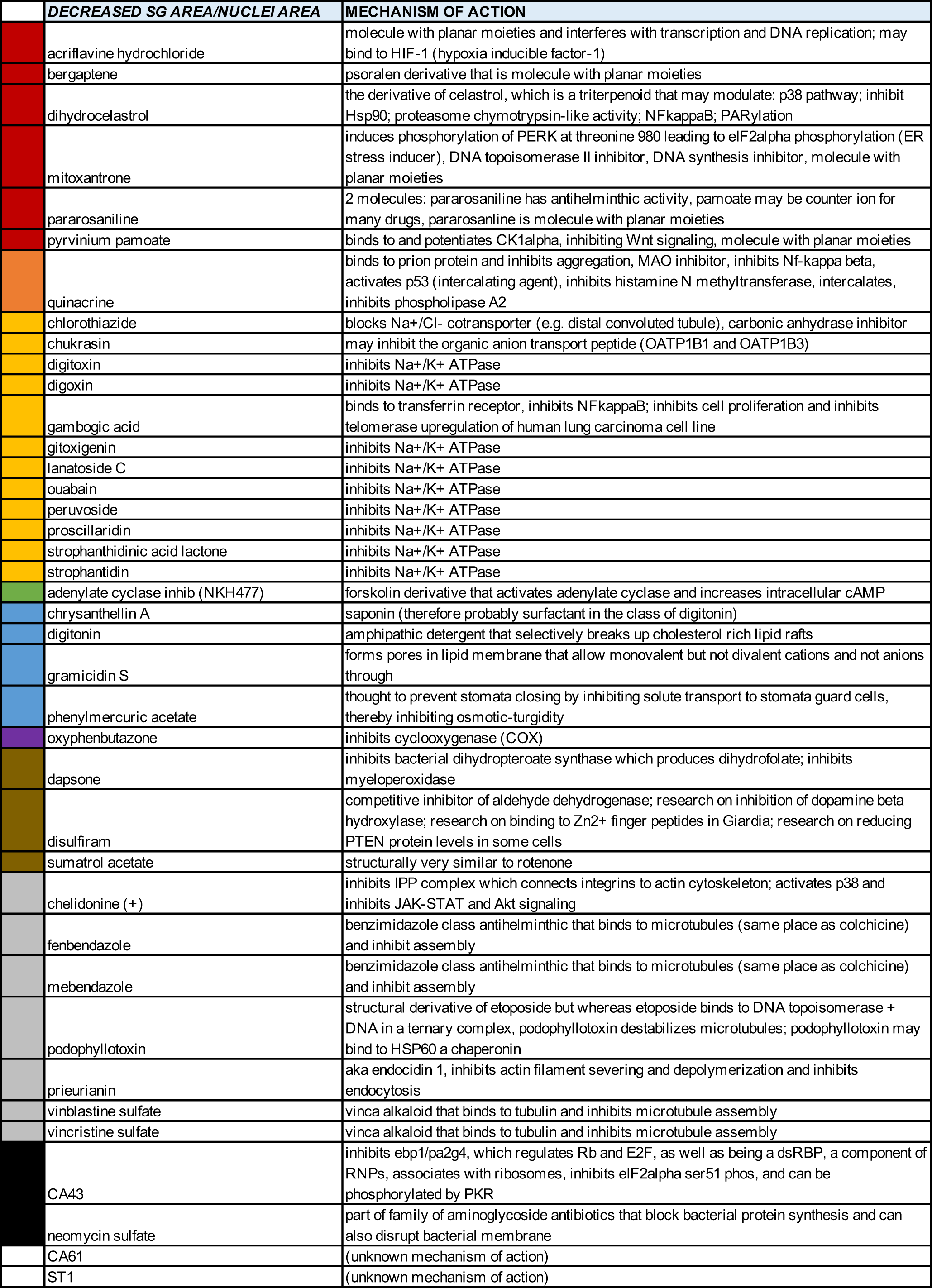
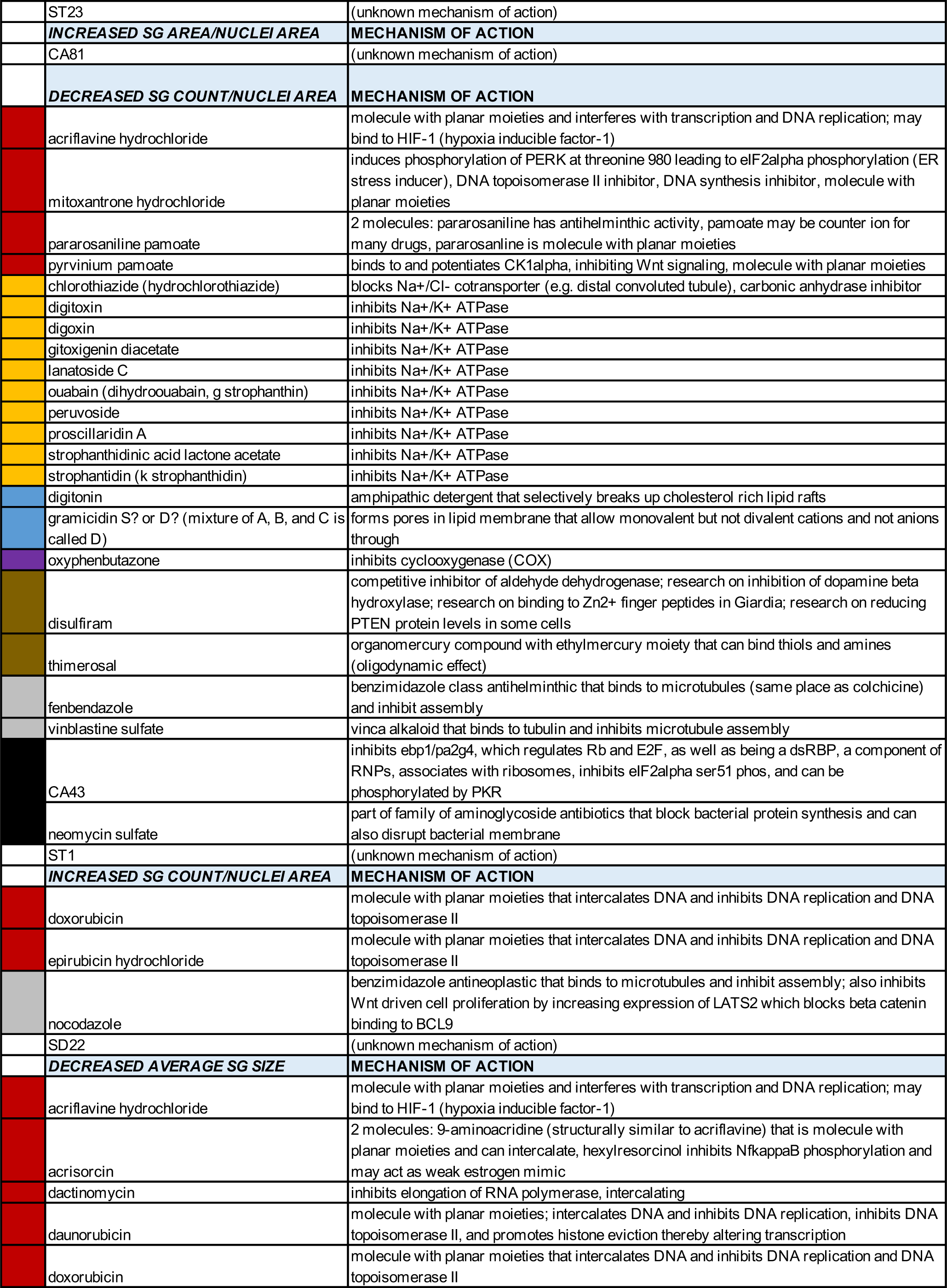
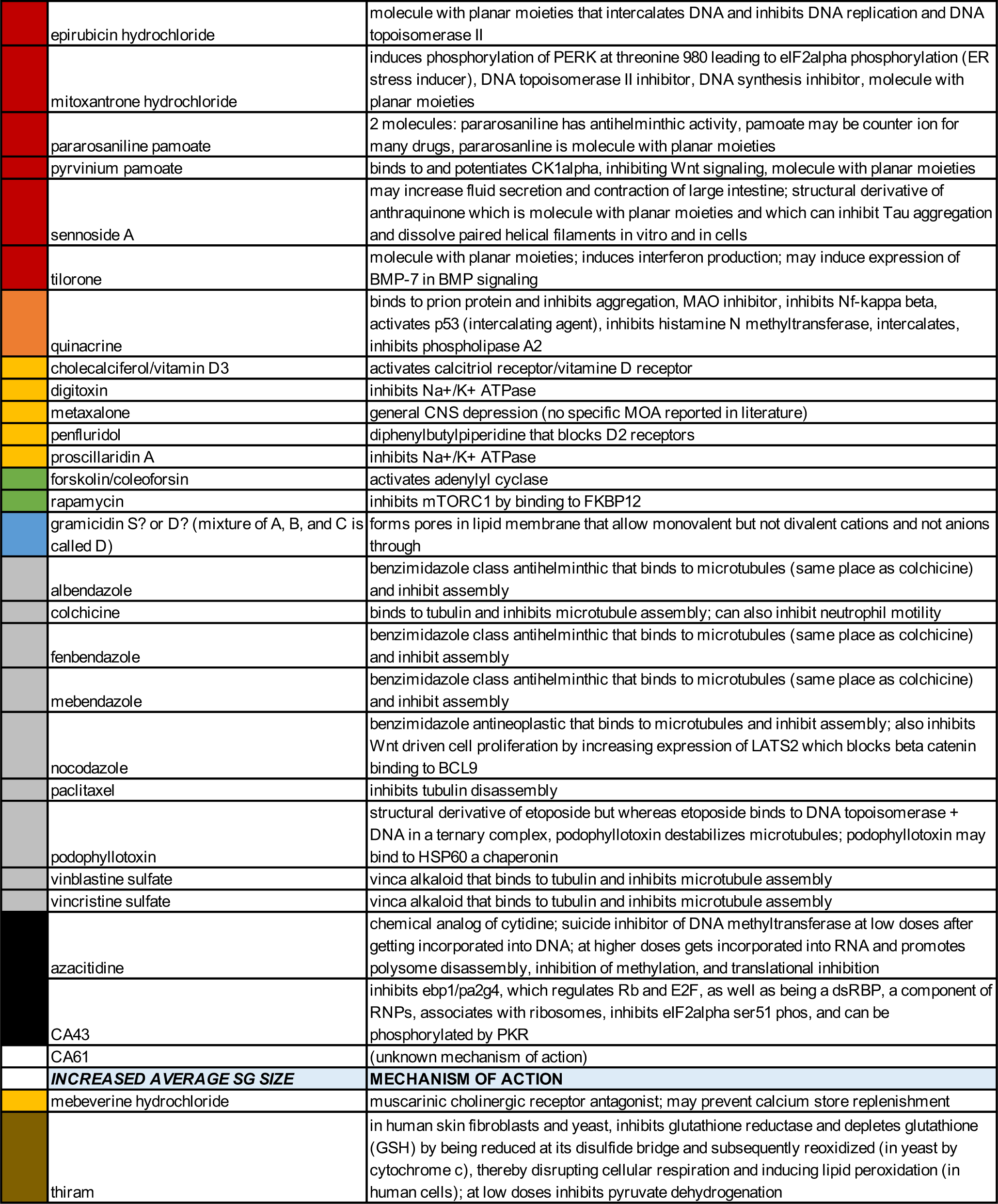
HEK293xT SG-modulating compounds and their annotated cellular targets. Table enumerating SG-modulating compounds identified in HEK293xT cells, color-coded by cellular target superclasses as in Table 1.

**Table 3.**
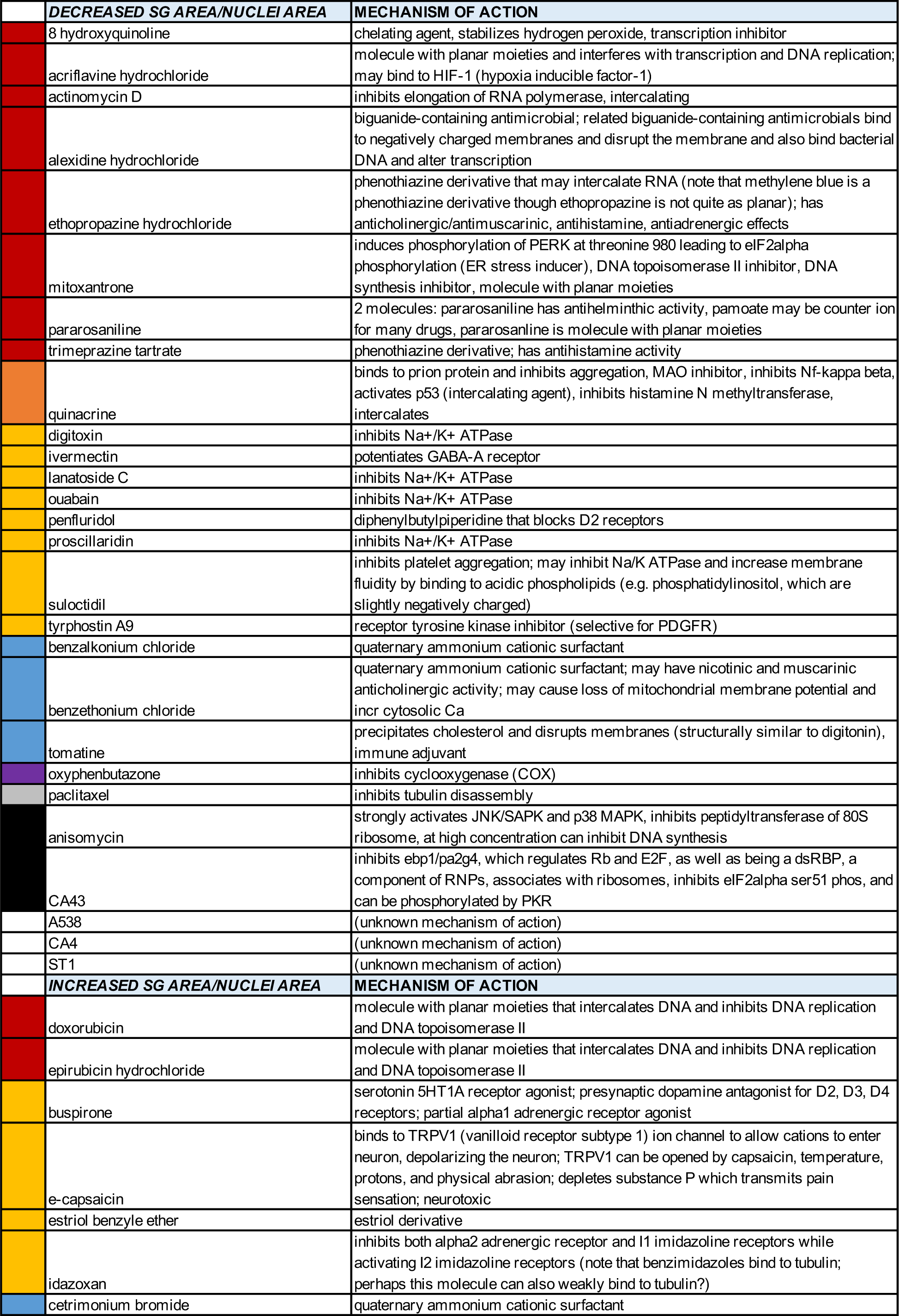
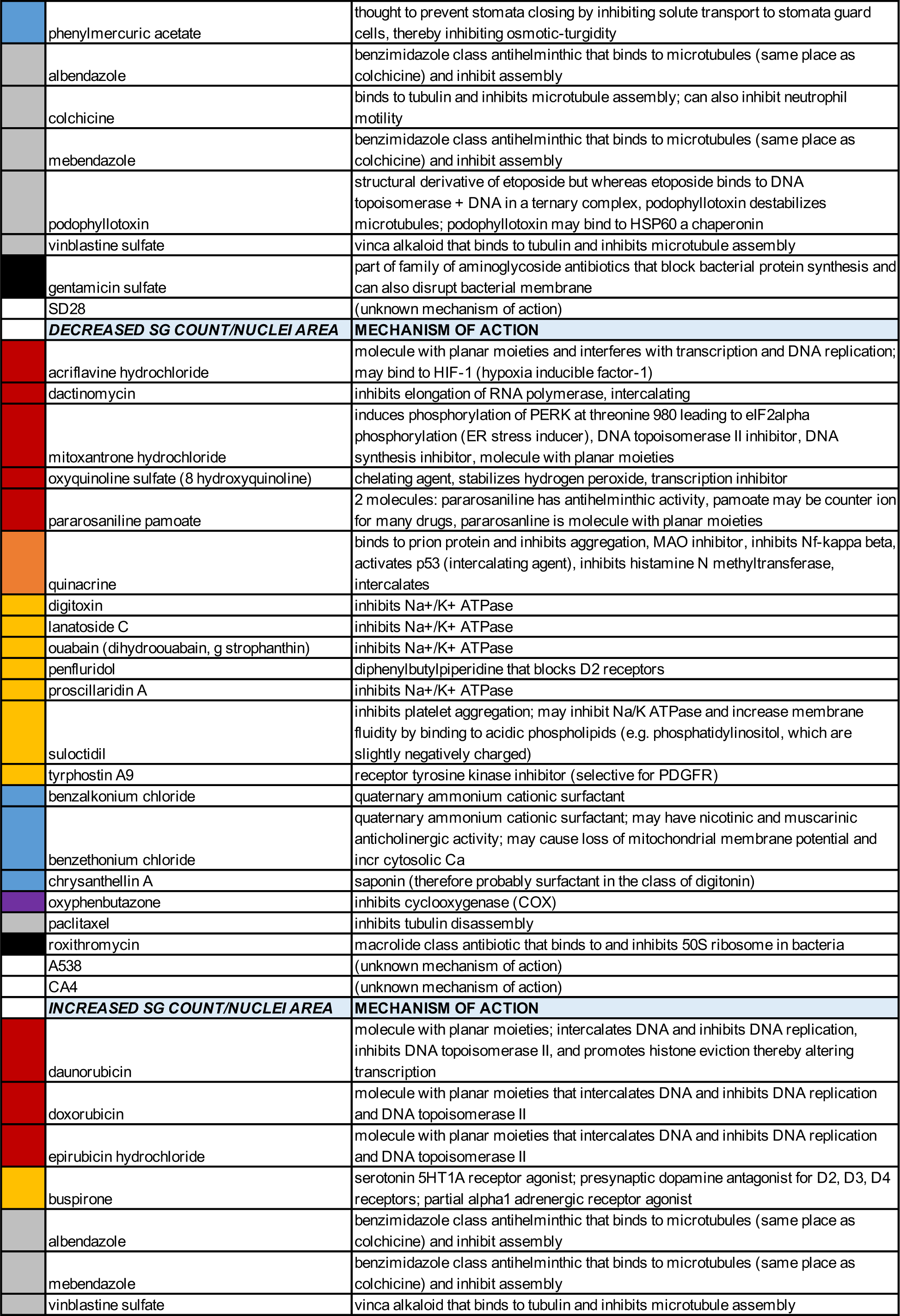
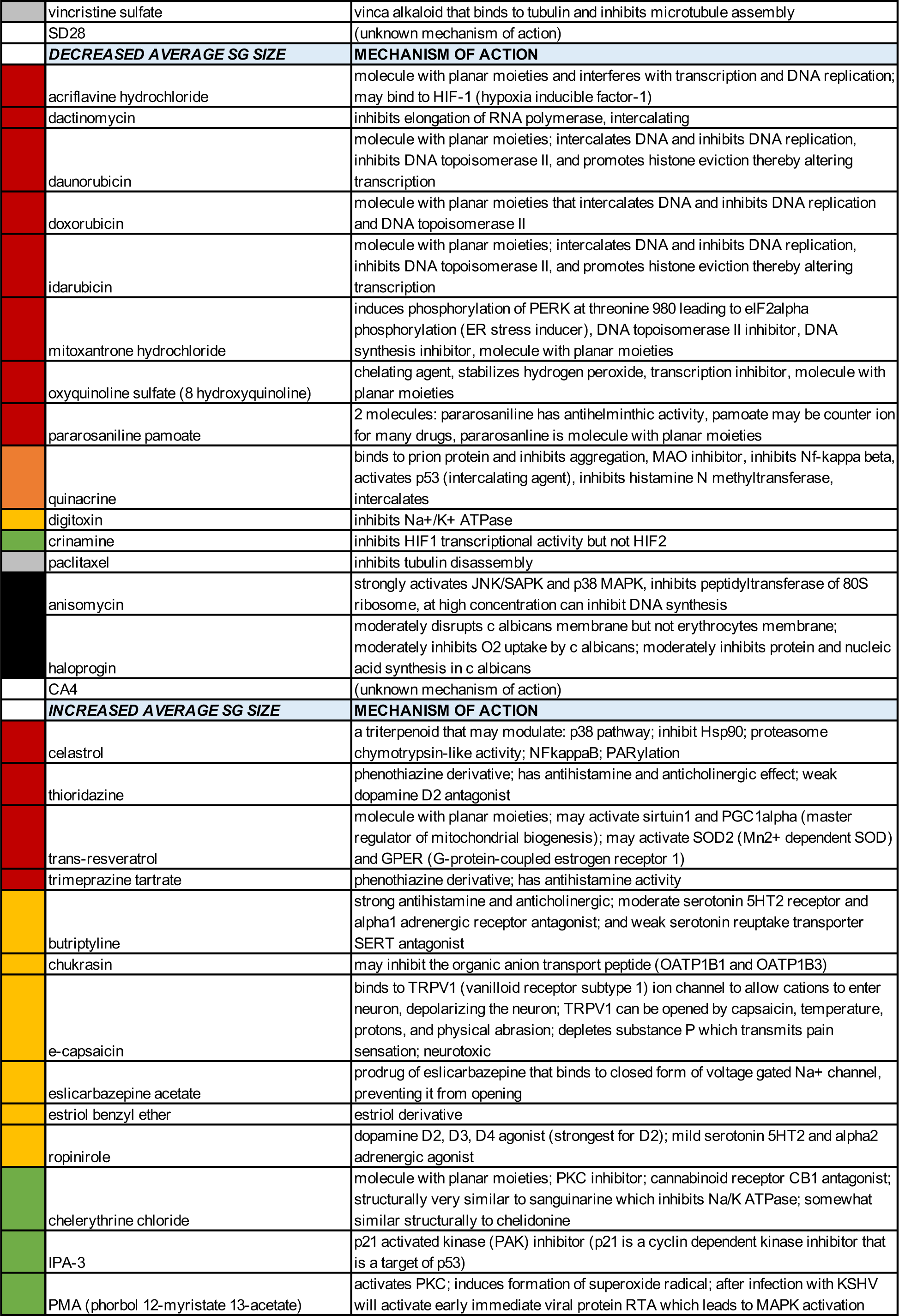
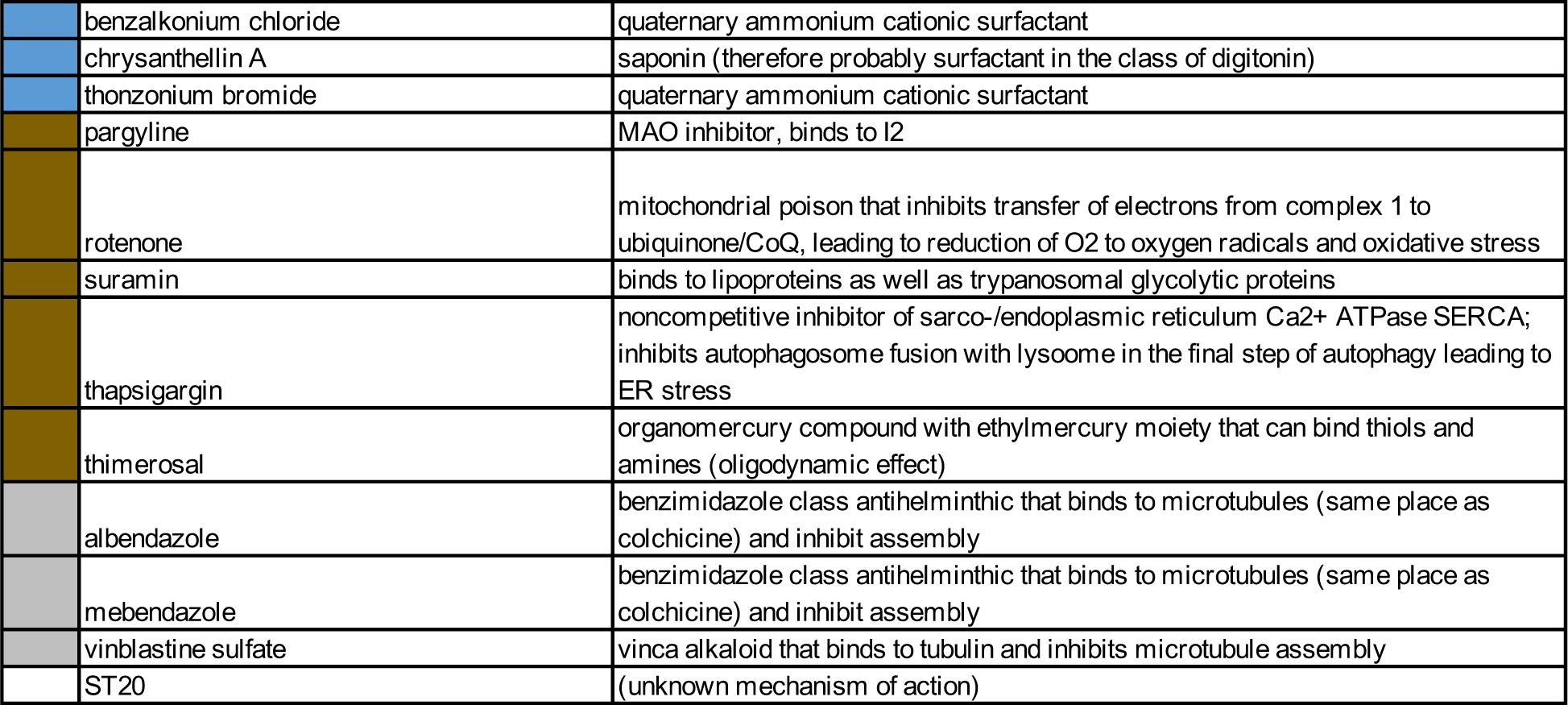
CV-B small molecule neural precursor cell SG-modulating compounds and their annotated cellular targets. Table enumerating SG-modulating compounds identified in CV-B small molecule neural precursor cells, color-coded by cellular target superclasses as in Table 1.

|  | DECREASED SG AREA/NUCLEI AREA | MECHANISM OF ACTION |
| --- | --- | --- |
|  | 8 hydroxyquinoline | chelating agent, stabilizes hydrogen peroxide, transcription inhibitor |
|  | acriflavine hydrochloride | molecule with planar moieties and interferes with transcription and DNA replication; may bind to HIF-1 (hypoxia inducible factor-1) |
|  | actinomycin D | inhibits elongation of RNA polymerase, intercalating |
|  | alexidine hydrochloride | biguanide-containing antimicrobial; related biguanide-containing antimicrobials bind to negatively charged membranes and disrupt the membrane and also bind bacterial DNA and alter transcription |
|  | ethopropazine hydrochloride | phenothiazine derivative that may intercalate RNA (note that methylene blue is a phenothiazine derivative though ethopropazine is not quite as planar); has anticholinergic/antimuscarinic, antihistamine, antiadrenergic effects |
|  | mitoxantrone | induces phosphorylation of PERK at threonine 980 leading to eIF2alpha phosphorylation (ER stress inducer), DNA topoisomerase II inhibitor, DNA synthesis inhibitor, molecule with planar moieties |
|  | pararosaniline | 2 molecules: pararosaniline has antihelminthic activity, pamoate may be counter ion for many drugs, pararosaniline is molecule with planar moieties |
|  | trimeprazine tartrate | phenothiazine derivative; has antihistamine activity |
|  | quinacrine | binds to prion protein and inhibits aggregation, MAO inhibitor, inhibits NF-kappa beta, activates p53 (intercalating agent), inhibits histamine N methyltransferase, intercalates |
|  | digitoxin | inhibits Na<sup>+</sup>/K<sup>+</sup> ATPase |
|  | ivermectin | potentiates GABA-A receptor |
|  | lanatoside C | inhibits Na<sup>+</sup>/K<sup>+</sup> ATPase |
|  | ouabain | inhibits Na<sup>+</sup>/K<sup>+</sup> ATPase |
|  | penfluridol | diphenylbutylpiperidine that blocks D2 receptors |
|  | proscillaridin | inhibits Na<sup>+</sup>/K<sup>+</sup> ATPase |
|  | suloctidil | inhibits platelet aggregation; may inhibit Na/K ATPase and increase membrane fluidity by binding to acidic phospholipids (e.g. phosphatidylinositol, which are slightly negatively charged) |
|  | tyrphostin A9 | receptor tyrosine kinase inhibitor (selective for PDGFR) |
|  | benzalkonium chloride | quaternary ammonium cationic surfactant |
|  | benzethonium chloride | quaternary ammonium cationic surfactant; may have nicotinic and muscarinic anticholinergic activity; may cause loss of mitochondrial membrane potential and incr cytosolic Ca |
|  | tomatine | precipitates cholesterol and disrupts membranes (structurally similar to digitonin), immune adjuvant |
|  | oxyphenbutazone | inhibits cyclooxygenase (COX) |
|  | paclitaxel | inhibits tubulin disassembly |
|  | anisomycin | strongly activates JNK/SAPK and p38 MAPK, inhibits peptidyltransferase of 80S ribosome, at high concentration can inhibit DNA synthesis |
|  | CA43 | inhibits ebp1/pa2g4, which regulates Rb and E2F, as well as being a dsRBP, a component of RNPs, associates with ribosomes, inhibits eIF2alpha ser51 phos, and can be phosphorylated by PKR |
|  | A538 | (unknown mechanism of action) |
|  | CA4 | (unknown mechanism of action) |
|  | ST1 | (unknown mechanism of action) |
|  | INCREASED SG AREA/NUCLEI AREA | MECHANISM OF ACTION |
|  | doxorubicin | molecule with planar moieties that intercalates DNA and inhibits DNA replication and DNA topoisomerase II |
|  | epirubicin hydrochloride | molecule with planar moieties that intercalates DNA and inhibits DNA replication and DNA topoisomerase II |
|  | buspirone | serotonin 5HT1A receptor agonist; presynaptic dopamine antagonist for D2, D3, D4 receptors; partial alpha1 adrenergic receptor agonist |
|  | e-capsaicin | binds to TRPV1 (vanilloid receptor subtype 1) ion channel to allow cations to enter neuron, depolarizing the neuron; TRPV1 can be opened by capsaicin, temperature, protons, and physical abrasion; depletes substance P which transmits pain sensation; neurotoxic |
|  | estriol benzyle ether | estriol derivative |
|  | idazoxan | inhibits both alpha2 adrenergic receptor and I1 imidazoline receptors while activating I2 imidazoline receptors (note that benzimidazoles bind to tubulin; perhaps this molecule can also weakly bind to tubulin?) |
|  | cetrimonium bromide | quaternary ammonium cationic surfactant |
|  | phenylmercuric acetate | thought to prevent stomata closing by inhibiting solute transport to stomata guard cells, thereby inhibiting osmotic-turgidity |
|  | albendazole | benzimidazole class antihelminthic that binds to microtubules (same place as colchicine) and inhibit assembly |
|  | colchicine | binds to tubulin and inhibits microtubule assembly; can also inhibit neutrophil motility |
|  | mebendazole | benzimidazole class antihelminthic that binds to microtubules (same place as colchicine) and inhibit assembly |
|  | podophyllotoxin | structural derivative of etoposide but whereas etoposide binds to DNA topoisomerase + DNA in a ternary complex, podophyllotoxin destabilizes microtubules; podophyllotoxin may bind to HSP60 a chaperonin |
|  | vinblastine sulfate | vinca alkaloid that binds to tubulin and inhibits microtubule assembly |
|  | gentamicin sulfate | part of family of aminoglycoside antibiotics that block bacterial protein synthesis and can also disrupt bacterial membrane |
|  | SD28 | (unknown mechanism of action) |
|  | **DECREASED SG COUNT/NUCLEI AREA** | **MECHANISM OF ACTION** |
|  | acriflavine hydrochloride | molecule with planar moieties and interferes with transcription and DNA replication; may bind to HIF-1 (hypoxia inducible factor-1) |
|  | dactinomycin | inhibits elongation of RNA polymerase, intercalating |
|  | mitoxantrone hydrochloride | induces phosphorylation of PERK at threonine 980 leading to eIF2alpha phosphorylation (ER stress inducer), DNA topoisomerase II inhibitor, DNA synthesis inhibitor, molecule with planar moieties |
|  | oxyquinoline sulfate (8 hydroxyquinoline) | chelating agent, stabilizes hydrogen peroxide, transcription inhibitor |
|  | pararosaniline pamoate | 2 molecules: pararosaniline has antihelminthic activity, pamoate may be counter ion for many drugs, pararosaniline is molecule with planar moieties |
|  | quinacrine | binds to prion protein and inhibits aggregation, MAO inhibitor, inhibits NF-kappa beta, activates p53 (intercalating agent), inhibits histamine N methyltransferase, intercalates |
|  | digitoxin | inhibits Na<sup>+</sup>/K<sup>+</sup> ATPase |
|  | lanatoside C | inhibits Na<sup>+</sup>/K<sup>+</sup> ATPase |
|  | ouabain (dihydroouabain, g strophanthin) | inhibits Na<sup>+</sup>/K<sup>+</sup> ATPase |
|  | penfluridol | diphenylbutylpiperidine that blocks D2 receptors |
|  | proscillaridin A | inhibits Na<sup>+</sup>/K<sup>+</sup> ATPase |
|  | suloctidil | inhibits platelet aggregation; may inhibit Na/K ATPase and increase membrane fluidity by binding to acidic phospholipids (e.g. phosphatidylinositol, which are slightly negatively charged) |
|  | tyrphostin A9 | receptor tyrosine kinase inhibitor (selective for PDGFR) |
|  | benzalkonium chloride | quaternary ammonium cationic surfactant |
|  | benzethonium chloride | quaternary ammonium cationic surfactant; may have nicotinic and muscarinic anticholinergic activity; may cause loss of mitochondrial membrane potential and incr cytosolic Ca |
|  | chrysanthellin A | saponin (therefore probably surfactant in the class of digitonin) |
|  | oxyphenbutazone | inhibits cyclooxygenase (COX) |
|  | paclitaxel | inhibits tubulin disassembly |
|  | roxithromycin | macrolide class antibiotic that binds to and inhibits 50S ribosome in bacteria |
|  | A538 | (unknown mechanism of action) |
|  | CA4 | (unknown mechanism of action) |
|  | **INCREASED SG COUNT/NUCLEI AREA** | **MECHANISM OF ACTION** |
|  | daunorubicin | molecule with planar moieties; intercalates DNA and inhibits DNA replication, inhibits DNA topoisomerase II, and promotes histone eviction thereby altering transcription |
|  | doxorubicin | molecule with planar moieties that intercalates DNA and inhibits DNA replication and DNA topoisomerase II |
|  | epirubicin hydrochloride | molecule with planar moieties that intercalates DNA and inhibits DNA replication and DNA topoisomerase II |
|  | buspirone | serotonin 5HT1A receptor agonist; presynaptic dopamine antagonist for D2, D3, D4 receptors; partial alpha1 adrenergic receptor agonist |
|  | albendazole | benzimidazole class antihelminthic that binds to microtubules (same place as colchicine) and inhibit assembly |
|  | mebendazole | benzimidazole class antihelminthic that binds to microtubules (same place as colchicine) and inhibit assembly |
|  | vinblastine sulfate | vinca alkaloid that binds to tubulin and inhibits microtubule assembly |
|  | vincristine sulfate | vinca alkaloid that binds to tubulin and inhibits microtubule assembly |
|  | SD28 | (unknown mechanism of action) |
|  | **DECREASED AVERAGE SG SIZE** | **MECHANISM OF ACTION** |
|  | acriflavine hydrochloride | molecule with planar moieties and interferes with transcription and DNA replication; may bind to HIF-1 (hypoxia inducible factor-1) |
|  | dactinomycin | inhibits elongation of RNA polymerase, intercalating |
|  | daunorubicin | molecule with planar moieties; intercalates DNA and inhibits DNA replication, inhibits DNA topoisomerase II, and promotes histone eviction thereby altering transcription |
|  | doxorubicin | molecule with planar moieties that intercalates DNA and inhibits DNA replication and DNA topoisomerase II |
|  | idarubicin | molecule with planar moieties; intercalates DNA and inhibits DNA replication, inhibits DNA topoisomerase II, and promotes histone eviction thereby altering transcription |
|  | mitoxantrone hydrochloride | induces phosphorylation of PERK at threonine 980 leading to eIF2alpha phosphorylation (ER stress inducer), DNA topoisomerase II inhibitor, DNA synthesis inhibitor, molecule with planar moieties |
|  | oxyquinoline sulfate (8 hydroxyquinoline) | chelating agent, stabilizes hydrogen peroxide, transcription inhibitor, molecule with planar moieties |
|  | pararosaniline pamoate | 2 molecules: pararosaniline has antihelminthic activity, pamoate may be counter ion for many drugs, pararosaniline is molecule with planar moieties |
|  | quinacrine | binds to prion protein and inhibits aggregation, MAO inhibitor, inhibits NF-kappa beta, activates p53 (intercalating agent), inhibits histamine N methyltransferase, intercalates |
|  | digitoxin | inhibits Na<sup>+</sup>/K<sup>+</sup> ATPase |
|  | crinamine | inhibits HIF1 transcriptional activity but not HIF2 |
|  | paclitaxel | inhibits tubulin disassembly |
|  | anisomycin | strongly activates JNK/SAPK and p38 MAPK, inhibits peptidyltransferase of 80S ribosome, at high concentration can inhibit DNA synthesis |
|  | haloprogin | moderately disrupts c albicans membrane but not erythrocytes membrane; moderately inhibits O2 uptake by c albicans; moderately inhibits protein and nucleic acid synthesis in c albicans |
|  | CA4 | (unknown mechanism of action) |
|  | **INCREASED AVERAGE SG SIZE** | **MECHANISM OF ACTION** |
|  | celastrol | a triterpenoid that may modulate: p38 pathway; inhibit Hsp90; proteasome chymotrypsin-like activity; NFkappaB; PARylation |
|  | thioridazine | phenothiazine derivative; has antihistamine and anticholinergic effect; weak dopamine D2 antagonist |
|  | trans-resveratrol | molecule with planar moieties; may activate sirtuin1 and PGC1alpha (master regulator of mitochondrial biogenesis); may activate SOD2 (Mn2+ dependent SOD) and GPER (G-protein-coupled estrogen receptor 1) |
|  | trimeprazine tartrate | phenothiazine derivative; has antihistamine activity |
|  | butriptyline | strong antihistamine and anticholinergic; moderate serotonin 5HT2 receptor and alpha1 adrenergic receptor antagonist; and weak serotonin reuptake transporter SERT antagonist |
|  | chukrasin | may inhibit the organic anion transport peptide (OATP1B1 and OATP1B3) |
|  | e-capsaicin | binds to TRPV1 (vanilloid receptor subtype 1) ion channel to allow cations to enter neuron, depolarizing the neuron; TRPV1 can be opened by capsaicin, temperature, protons, and physical abrasion; depletes substance P which transmits pain sensation; neurotoxic |
|  | eslicarbazepine acetate | prodrug of eslicarbazepine that binds to closed form of voltage gated Na<sup>+</sup> channel, preventing it from opening |
|  | estriol benzyl ether | estriol derivative |
|  | ropinirole | dopamine D2, D3, D4 agonist (strongest for D2); mild serotonin 5HT2 and alpha2 adrenergic agonist |
|  | chelerythrine chloride | molecule with planar moieties; PKC inhibitor; cannabinoid receptor CB1 antagonist; structurally very similar to sanguinarine which inhibits Na/K ATPase; somewhat similar structurally to chelidonine |
|  | IPA-3 | p21 activated kinase (PAK) inhibitor (p21 is a cyclin dependent kinase inhibitor that is a target of p53) |
|  | PMA (phorbol 12-myristate 13-acetate) | activates PKC; induces formation of superoxide radical; after infection with KSHV will activate early immediate viral protein RTA which leads to MAPK activation |

|  |  |
| --- | --- |
| benzalkonium chloride | quatarnary ammonium cationic surfactant |
| chrysanthellin A | saponin (therefore probably surfactant in the class of digitonin) |
| thonzonium bromide | quatarnary ammonium cationic surfactant |
| pargyline | MAO inhibitor, binds to I2 |
| rotenone | mitochondrial poison that inhibits transfer of electrons from complex 1 to ubiquinone/CoQ, leading to reduction of O2 to oxygen radicals and oxidative stress |
| suramin | binds to lipoproteins as well as trypanosomal glycolytic proteins |
| thapsigargin | noncompetitive inhibitor of sarco-/endoplasmic reticulum Ca2+ ATPase SERCA; inhibits autophagosome fusion with lysosome in the final step of autophagy leading to ER stress |
| thimerosal | organomercury compound with ethylmercury moiety that can bind thiols and amines (oligodynamic effect) |
| albendazole | benzimidazole class antihelminthic that binds to microtubules (same place as colchicine) and inhibit assembly |
| mebendazole | benzimidazole class antihelminthic that binds to microtubules (same place as colchicine) and inhibit assembly |
| vinblastine sulfate | vinca alkaloid that binds to tubulin and inhibits microtubule assembly |
| ST20 | (unknown mechanism of action) |

**Table 4.** HEK293xT and CV-B small molecule neural precursor cell overlapping SG-modulating compounds and their annotated cellular targets. Table enumerating SG-modulating compounds identified in both HEK293xT cells and CV-B small molecule neural precursor cells, color-coded by cellular target superclasses as in Table 1.

|  | DECREASED SG AREA/NUCLEI AREA | MECHANISM OF ACTION |
| --- | --- | --- |
|  | acriflavine hydrochloride | molecule with planar moieties and interferes with transcription and DNA replication; may bind to HIF-1 (hypoxia inducible factor-1) |
|  | mitoxantrone | induces phosphorylation of PERK at threonine 980 leading to eIF2alpha phosphorylation (ER stress inducer), DNA topoisomerase II inhibitor, DNA synthesis inhibitor, molecule with planar moieties |
|  | pararosaniline | 2 molecules: pararosaniline has antihelminthic activity, pamoate may be counter ion for many drugs, pararosaniline is molecule with planar moieties |
|  | quinacrine | binds to prion protein and inhibits aggregation, MAO inhibitor, inhibits Nf-kappa beta, activates p53 (intercalating agent), inhibits histamine N methyltransferase, intercalates, inhibits phospholipase A2 |
|  | digitoxin | inhibits Na+/K+ ATPase |
|  | lanatoside C | inhibits Na+/K+ ATPase |
|  | ouabain | inhibits Na+/K+ ATPase |
|  | proscillaridin | inhibits Na+/K+ ATPase |
|  | oxyphenbutazone | inhibits cyclooxygenase (COX) |
|  | CA43 | inhibits ebp1/pa2g4, which regulates Rb and E2F, as well as being a dsRBP, a component of RNPs, associates with ribosomes, inhibits eIF2alpha ser51 phos, and can be phosphorylated by PKR |
|  | ST1 | (unknown mechanism of action) |
|  | DECREASED SG COUNT/NUCLEI AREA | MECHANISM OF ACTION |
|  | acriflavine hydrochloride | molecule with planar moieties and interferes with transcription and DNA replication; may bind to HIF-1 (hypoxia inducible factor-1) |
|  | mitoxantrone hydrochloride | induces phosphorylation of PERK at threonine 980 leading to eIF2alpha phosphorylation (ER stress inducer), DNA topoisomerase II inhibitor, DNA synthesis inhibitor, molecule with planar moieties |
|  | pararosaniline pamoate | 2 molecules: pararosaniline has antihelminthic activity, pamoate may be counter ion for many drugs, pararosaniline is molecule with planar moieties |
|  | digitoxin | inhibits Na+/K+ ATPase |
|  | lanatoside C | inhibits Na+/K+ ATPase |
|  | ouabain (dihydroouabain, g strophanthin) | inhibits Na+/K+ ATPase |
|  | proscillaridin A | inhibits Na+/K+ ATPase |
|  | oxyphenbutazone | inhibits cyclooxygenase (COX) |
|  | INCREASED SG COUNT/NUCLEI AREA | MECHANISM OF ACTION |
|  | doxorubicin | molecule with planar moieties that intercalates DNA and inhibits DNA replication and DNA topoisomerase II |
|  | epirubicin hydrochloride | molecule with planar moieties that intercalates DNA and inhibits DNA replication and DNA topoisomerase II |
|  | DECREASED AVERAGE SG SIZE | MECHANISM OF ACTION |
|  | acriflavine hydrochloride | molecule with planar moieties and interferes with transcription and DNA replication; may bind to HIF-1 (hypoxia inducible factor-1) |
|  | dactinomycin | inhibits elongation of RNA polymerase, intercalating |
|  | daunorubicin | molecule with planar moieties; intercalates DNA and inhibits DNA replication, inhibits DNA topoisomerase II, and promotes histone eviction thereby altering transcription |
|  | doxorubicin | molecule with planar moieties that intercalates DNA and inhibits DNA replication and DNA topoisomerase II |
|  | mitoxantrone hydrochloride | induces phosphorylation of PERK at threonine 980 leading to eIF2alpha phosphorylation (ER stress inducer), DNA topoisomerase II inhibitor, DNA synthesis inhibitor, molecule with planar moieties |
|  | pararosaniline pamoate | 2 molecules: pararosaniline has antihelminthic activity, pamoate may be counter ion for many drugs, pararosaniline is molecule with planar moieties |
|  | quinacrine | binds to prion protein and inhibits aggregation, MAO inhibitor, inhibits Nf-kappa beta, activates p53 (intercalating agent), inhibits histamine N methyltransferase, intercalates, inhibits phospholipase A2 |
|  | digitoxin | inhibits Na+/K+ ATPase |
|  | paclitaxel | inhibits tubulin disassembly |

Among our hit compounds which have not previously been reported to modulate SG formation, we identified many that act on targets at the surface of cells, such as ion channels, surface receptors, transporters, and lipid membranes (Figures 2E, S2A, and Tables 2–4). Modulation of these plasma membrane targets has not previously been reported to directly modulate SGs. We identified a large group of cardiac glycosides including digitoxin and proscillaridin A which inhibit the Na^+^/K^+^-ATPase, as well as the GABA-A chloride channel agonist ivermectin, compounds that bind to voltage gated Na^+^ channels such as eslicarbazepine, and amphipathic molecules such as benzethonium which can disrupt lipid membranes (Figure 2C and Tables 2–4). Intriguingly, cardiac glycosides were previously shown to disperse nuclear TDP-43 foci in ALS patient-derived iPS-MNs, though it remains unclear how these compounds exert this effect and whether these nuclear TDP-43 foci are related to SGs (*39*). We also identified the receptor tyrosine kinase inhibitor tyrphostin A9 as a SG-modulating compound, which was reported in a prior study as increasing the survival of MNs carrying SOD1 mutations; however it is uncertain whether this survival effect is related to the ability of tyrphostin A9 to reduce the amount of SG formation per cell (*40*). Interestingly, the D2 receptor antagonist penfluridol reduced the amount of SG formation per cell in NPCs under NaAsO_2_ stress but inversely modestly induced SG formation in the absence of stress (Figure S2C). Overall, it is possible that several of these compounds which act on targets at the surface of cells modulate SGs by interfering with intracellular ion, solute, and/or osmolarity homeostasis. It has been reported that physical-chemical properties such as ionic content, tonicity, and concentration of hydrotrope solutes such as ATP regulate liquid-liquid phase transitions, which may be important for SG formation (*41, 42*). The divalent cation Zn^2+^ has also been recently reported to regulate the phase transition of the SG-associated protein TIA1 (*43*).

We also identified SG-modulating compounds that target a broad range of intracellular processes, including signaling pathways and transcription factors, metabolism, and inflammatory molecule biosynthesis pathways (Figures 2E, S2A and Tables 2–4). Several compounds modulate intracellular signaling pathways, including two compounds which target DYRK and RACK1, proteins at the intersection of SGs/mTOR signaling and SGs/SAPK signaling, respectively (*7, 8*). Some compounds modulate intracellular metabolic processes which are known to regulate SGs, including heat shock proteins, autophagosomes, and proteasomes (*44–46*). We identified seven hit compounds that have anti-inflammatory activities, including quinacrine, an inhibitor of phospholipase A2 (PLA2), and oxyphenbutazone, a cyclooxygenase inhibitor (Figure 2C and Tables 2–4).

Strikingly, we identified a large group of SG modulatory compounds that contain extended planar molecular moieties, many of which act as nucleic acid intercalating molecules, including mitoxantrone, doxorubicin, and daunorubicin (Figures 2E, S2A, and Tables 2–4). Molecules with planar moieties and nucleic acid intercalating compounds have not previously been reported to modulate SGs, but these compounds strongly reduced the amount of SG formation per cell, altered the average SG size, and/or changed the number of SGs per cell (Figure 2C and Tables 2–4). Interestingly, doxorubicin has recently been reported to modulate the phase transition diagram of RNA-only condensates containing CAG or CUG repeats *in vitro* and *in vivo* (*3, 47*). Furthermore, quinacrine, another molecule with planar moieties, has previously been reported to directly bind to the prion-like domain of the prion protein PrP and inhibit aggregation *in vitro* (*48*). It is possible these molecules with planar moieties modulate SGs by interacting with RNAs or protein IDRs contained therein.

### Screen Hit Compounds Robustly Inhibit SG Formation Across Different Stress Contexts

We decided to evaluate the two most highly represented groups of hit compounds – molecules with planar moieties and cardiac glycosides. As it is unclear how these compounds modulate SGs, investigations with these compounds are likely to reveal molecular rules underlying SG biology. First, we tested 16 and 18 compounds (Tables 5–6) in their capacities to reduce the amount of SG formation per cell in HEK293xT cells and NPCs, respectively, in a dose dependent manner over a concentration range covering three orders of magnitude and spanning the initial screening concentration of 10 μM (Figure 3A). We found that 11 and 12 compounds show concentration dependent inhibition of SG formation in HEK293xT cells and NPCs, respectively (Figures 3B and S3A). After fitting logistic functions to the dose response data, we estimated the compounds’ 50% inhibitory concentrations (IC50s; Figure 3B). Most compounds’ IC50s were in the single digit micromolar range, which suggests that their mechanisms of actions are unlikely a direct inhibition of one specific protein or target, but rather the sum of activities against several targets.

**Figure 3.**
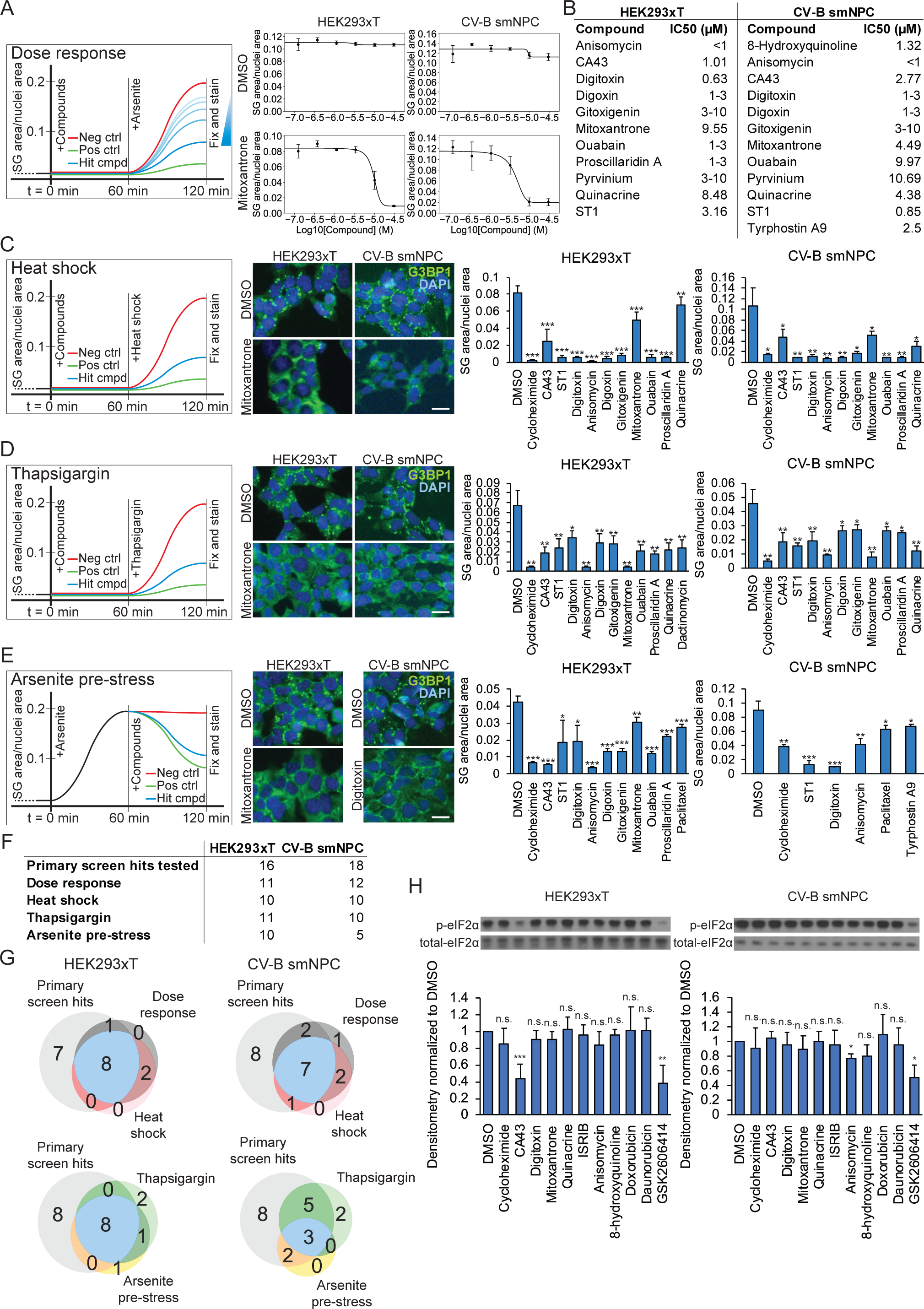
Counterscreen validation of SG inhibiting compounds. (A) Overview of the assay used to demonstrate dose-dependent inhibition of SG formation by SG screen hit compounds. Scatterplot and logistic regression showing dose-dependent reduction of SG area/nuclei area by a nucleic acid intercalating molecule mitoxantrone but not by the vehicle control DMSO. Error bars are sample standard deviations from four biological replicates. SG: stress granule, NPC: small molecule neural precursor cells. (B) Table summarizing the estimated IC50s of SG inhibiting compounds, determined from the midpoints of logistic regression functions fitted to dose response scatterplots. (C) Overview of the assay used to test the efficacy of compounds to inhibit SG formation under heat shock stress. Representative wide-field fluorescence microscopy of HEK293xT cells and CV-B small molecule neural precursor cells treated with a nucleic acid intercalating molecule mitoxantrone versus DMSO control under heat shock stress. Histograms showing the reduction in SG area/nuclei area in HEK293xT cells and CV-B small molecule neural precursor cells treated with hit compounds under heat shock stress. (D) Overview of the assay used to test the efficacy of compounds to inhibit SG formation under thapsigargin stress. Representative wide-field fluorescence microscopy of HEK293xT cells and CV-B small molecule neural precursor cells treated with a nucleic acid intercalating molecule mitoxantrone versus DMSO control under thapsigargin stress. Histograms showing the reduction in SG area/nuclei area in HEK293xT cells and CV-B small molecule neural precursor cells treated with hit compounds under thapsigargin stress. (E) Overview of the assay used to test the efficacy of compounds to reverse SG formation after cells have been pre-stressed by NaAsO_2_ for one hour. Representative wide-field fluorescence microscopy of HEK293xT cells and CV-B small molecule neural precursor cells treated with a nucleic acid intercalating molecule mitoxantrone or Na^+^/K^+^-ATPase inhibitor digitoxin versus DMSO control after one hour of NaAsO_2_ pre-stress. Histograms showing the reduction in SG area/nuclei area in HEK293xT cells and CV-B small molecule neural precursor cells when hit compounds are added after NaAsO_2_ stress. (C-E) Significance levels are from two-tailed two sample Student’s t-tests to DMSO control: * p < 0.05, ** p < 0.01, *** p < 0.001. Error bars are sample standard deviations from four biological replicates. Scale bar is 50 μm. (F) Table summarizing the number of compounds that reduced SG area/nuclei area in each of the counterscreen assays. (G) Venn diagrams representing the number of compounds that reduced SG area/nuclei area in more than one counterscreen assay. (H) Representative Western blots of eIF2α phosphorylated at serine 51 versus total eIF2α protein in cells treated with hit compounds and under NaAsO_2_ stress. Histograms quantifying the amount of eIF2α protein phosphorylated at serine 51 in HEK293xT cells and CV-B small molecule neural precursor cells treated with SG inhibiting compounds under NaAsO_2_ stress. Histogram values are normalized to samples treated with the vehicle control DMSO. Significance levels are from two-tailed one sample Student’s t-tests: * p < 0.05, ** p < 0.01, *** p < 0.001. Error bars are sample standard deviations from at least four biological replicates.

**Table 5.** Screen hit compounds tested in HEK293xT cell counterscreens. Table enumerating SG screen hit compounds that were tested in HEK293xT cells in four counterscreen assays: dose response assay against NaAsO_2_ stress, heat shock stress assay, thapsigargin stress assay, and NaAsO_2_ pre-stress assay. Compounds color-coded by cellular target superclasses as in Table 1.

|  | COMPOUND | MECHANISM OF ACTION |
| --- | --- | --- |
|  | mitoxantrone | induces phosphorylation of PERK at threonine 980 leading to eIF2alpha phosphorylation (ER stress inducer), DNA topoisomerase II inhibitor, DNA synthesis inhibitor, molecule with planar moieties |
|  | pararosaniline | 2 molecules: pararosaniline has antihelminthic activity, pamoate may be counter ion for many drugs, pararosaniline is molecule with planar moieties |
|  | pyrvinium pamoate | binds to and potentiates CK1alpha, inhibiting Wnt signaling, molecule with planar moieties |
|  | chlorothiazide<br>(hydrochlorothiazide) | blocks Na<sup>+</sup>/Cl<sup>-</sup> cotransporter (e.g. distal convoluted tubule), carbonic anhydrase inhibitor |
|  | digitoxin | inhibits Na<sup>+</sup>/K<sup>+</sup> ATPase |
|  | digoxin | inhibits Na<sup>+</sup>/K<sup>+</sup> ATPase |
|  | gitoxigenin | inhibits Na<sup>+</sup>/K<sup>+</sup> ATPase |
|  | lanatoside C | inhibits Na<sup>+</sup>/K<sup>+</sup> ATPase |
|  | ouabain | inhibits Na<sup>+</sup>/K<sup>+</sup> ATPase |
|  | peruvoside | inhibits Na<sup>+</sup>/K<sup>+</sup> ATPase |
|  | proscillaridin | inhibits Na<sup>+</sup>/K<sup>+</sup> ATPase |
|  | strophanthidinic acid lactone | inhibits Na<sup>+</sup>/K<sup>+</sup> ATPase |
|  | strophantidin | inhibits Na<sup>+</sup>/K<sup>+</sup> ATPase |
|  | gramicidin S | forms pores in lipid membrane that allow monovalent but not divalent cations and not anions through |
|  | CA43 | inhibits ebp1/pa2g4, which regulates Rb and E2F, as well as being a dsRBP, a component of RNPs, associates with ribosomes, inhibits eIF2alpha ser51 phos, and can be phosphorylated by PKR |
|  | ST1 | (unknown mechanism of action) |

**Table 6.** Screen hit compounds tested in CV-B small molecule neural precursor cell counterscreens. Table enumerating SG screen hit compounds that were tested in NPCs in four counterscreen assays: dose response assay against NaAsO_2_, heat shock and thapsigargin stress assays as well as NaAsO_2_ pre-stress assay. Compounds color-coded by cellular target superclasses as in Table 1.

|  | COMPOUND | MECHANISM OF ACTION |
| --- | --- | --- |
|  | 8 hydroxyquinoline | chelating agent, stabilizes hydrogen peroxide, transcription inhibitor |
|  | dactinomycin | inhibits elongation of RNA polymerase, intercalating |
|  | mitoxantrone | induces phosphorylation of PERK at threonine 980 leading to eIF2alpha phosphorylation (ER stress inducer), DNA topoisomerase II inhibitor, DNA synthesis inhibitor, molecule with planar moieties |
|  | pararosaniline | 2 molecules: pararosaniline has antihelminthic activity, pamoate may be counter ion for many drugs, pararosaniline is molecule with planar moieties |
|  | quinacrine | binds to prion protein and inhibits aggregation, MAO inhibitor, inhibits Nf-kappa beta, activates p53 (intercalating agent), inhibits histamine N methyltransferase, intercalates |
|  | digitoxin | inhibits Na<sup>+</sup>/K<sup>+</sup> ATPase |
|  | ivermectin | potentiates GABA-A receptor |
|  | lanatoside C | inhibits Na<sup>+</sup>/K<sup>+</sup> ATPase |
|  | ouabain | inhibits Na<sup>+</sup>/K<sup>+</sup> ATPase |
|  | penfluridol | diphenylbutylpiperidine that blocks D2 receptors |
|  | proscillaridin | inhibits Na<sup>+</sup>/K<sup>+</sup> ATPase |
|  | tyrphostin A9 | receptor tyrosine kinase inhibitor (selective for PDGFR) |
|  | tomatine | precipitates cholesterol and disrupts membranes (structurally similar to digitonin), immune adjuvant |
|  | oxyphenbutazone | inhibits cyclooxygenase (COX) |
|  | paclitaxel | inhibits tubulin disassembly |
|  | anisomycin | strongly activates JNK/SAPK and p38 MAPK, inhibits peptidyltransferase of 80S ribosome, at high concentration can inhibit DNA synthesis |
|  | CA43 | inhibits ebp1/pa2g4, which regulates Rb and E2F, as well as being a dsRBP, a component of RNPs, associates with ribosomes, inhibits eIF2alpha ser51 phos, and can be phosphorylated by PKR |
|  | ST1 | (unknown mechanism of action) |

NaAsO_2_ generates intracellular oxidative damage that leads to activation of the heme-regulated inhibitor (HRI) kinase, which phosphorylates eIF2α leading to SG assembly. On the other hand, heat shock and thapsigargin-induced endoplasmic reticulum stress stimulate the unfolded protein response, resulting in activation of protein kinase RNA-like endoplasmic reticulum kinase (PERK), which phosphorylates eIF2α (*2*). Thapsigargin also inhibits the autophagosome, hindering SG clearance through autophagy (*44*). We therefore tested whether the hit compounds inhibit SG formation induced by heat shock or the autophagosome and endoplasmic reticulum (ER) poison thapsigargin. We found that at least 10 compounds significantly reduced the amount of SG formation per cell under heat shock or thapsigargin poisoning in both HEK293xT cells and NPCs (Figures 3C-D). These alternate stressors induce SG formation via signaling through orthogonal cellular stress pathways compared to NaAsO_2_ (*2*). Because the hit compounds inhibited SG formation under these orthogonal stressors, the mechanisms of action for these compounds are broader than just antioxidants scavenging reactive radicals generated by NaAsO_2_ or inhibitors of HRI kinase.

Lastly, we tested whether these hit compounds can disassemble pre-existing SGs, by pre-stressing cells with NaAsO_2_ and then adding compounds after SGs have already developed. We identified 10 and 5 compounds which significantly disassembled pre-formed SGs in HEK293xT cells and NPCs, respectively (Figure 3E). This confirms that our screening paradigm of adding compounds first and stressing second successfully identified compounds which prevent SG formation, but also compounds which induced disassembly of SGs after they have already begun to form. In summary, the compounds tested in the counterscreens, most of which are molecules with planar moieties or cardiac glycosides, robustly inhibited SGs across diverse stress conditions (Figures 3F-G).

### Hit Compounds Inhibit SG formation in a Manner Independent of eIF2α Phosphorylation

Next, we evaluated if our compounds that inhibit SG formation affected eIF2α phosphorylation or levels. Unexpectedly, most of the hit compounds did not alter NaAsO_2_-induced phosphorylation of eIF2α at serine 51 or total levels of eIF2α, whereas our control GSK2606414, a selective PERK inhibitor, significantly reduced eIF2α phosphorylation as previously reported (Figures 3H and S3C) (*34*). Oddly, in the absence of stress, the EBP1 kinase modulating compound CA43, the cardiac glycoside digitoxin, the ribosomal inhibitor anisomycin, and the planar heterocyclic molecule 8-hydroxyquinoline significantly increased phosphorylation of eIF2α above baseline without changing total levels of eIF2α, as did the PERK inhibitor GSK2606414, all without inducing SG formation (Figures S2C, S3B, and S3D). These results indicate that these hit compounds likely act on SGs themselves rather than on upstream kinases or signaling pathways that lead to phosphorylation of eIF2α.

### Screen Compounds Inhibit SG Formation in iPS-MNs

As our primary and counterscreens were performed in proliferative cells, we wanted to evaluate if the hit compounds can also inhibit SG formation in iPS-MNs terminally differentiated from iPSC lines. We utilized 4 iPSC lines from patients carrying ALS-linked mutations in the C-terminal IDR of TDP-43 (N352S and G298S), 2 iPSC lines from patients harboring mutations in the C-terminal NLS of FUS (R521G), and 4 control iPSC lines from healthy controls who are genetically related to the individuals carrying ALS-associated mutations (Figure S4A). We selected 10 hit compounds that were validated as SG inhibiting compounds in the HEK293xT cell and NPC counterscreens and were also representative molecules spanning most of the hit compound classes identified in the screen, including several molecules with planar moieties (Table 7). We also included the positive control cycloheximide and the negative control DMSO (Table 7).

**Table 7.** Screen hit compounds tested for SG inhibition in iPS-MNs. Table enumerating SG screen hit compounds that were tested in control, *TARDBP* mutant, or *FUS* mutant MNs against NaAsO_2_ stress, thapsigargin stress, or puromycin stress. Compounds color-coded by cellular target superclasses as in Table 1.

|  | COMPOUND | MECHANISM OF ACTION |
| --- | --- | --- |
|  | DMSO | negative control; vehicle |
|  | cycloheximide | positive control; inhibits ribosomal translocation |
|  | 8 hydroxyquinoline | chelating agent, stabilizes hydrogen peroxide, transcription inhibitor |
|  | daunorubicin | molecule with planar moieties; intercalates DNA and inhibits DNA replication, inhibits DNA topoisomerase II, and promotes histone eviction thereby altering transcription |
|  | mitoxantrone | induces phosphorylation of PERK at threonine 980 leading to eIF2alpha phosphorylation (ER stress inducer), DNA topoisomerase II inhibitor, DNA synthesis inhibitor, molecule with planar moieties |
|  | pyrvinium pamoate | binds to and potentiates CK1alpha, inhibiting Wnt signaling, molecule with planar moieties |
|  | quinacrine | binds to prion protein and inhibits aggregation, MAO inhibitor, inhibits Nf-kappa beta, activates p53 (intercalating agent), inhibits histamine N methyltransferase, intercalates |
|  | digitoxin | inhibits Na<sup>+</sup>/K<sup>+</sup> ATPase |
|  | penfluridol | diphenylbutylpiperidine that blocks D2 receptors |
|  | benzethonium chloride | quaternary ammonium cationic surfactant; may have nicotinic and muscarinic anticholinergic activity; may cause loss of mitochondrial membrane potential and incr cytosolic Ca |
|  | anisomycin | strongly activates JNK/SAPK and p38 MAPK, inhibits peptidyltransferase of 80S ribosome, at high concentration can inhibit DNA synthesis |
|  | CA43 | inhibits ebp1/pa2g4, which regulates Rb and E2F, as well as being a dsRBP, a component of RNPs, associates with ribosomes, inhibits eIF2alpha ser51 phos, and can be phosphorylated by PKR |

Puromycin which has previously been shown to robustly induce SG formation in MNs (*1, 11, 23, 45*) was utilized in our post-mitotic iPS-MNs. We observed that our compounds were effective at inhibiting SG formation across all lines, regardless of mutation status (Figures 4A-C). Six (of 10) compounds – CA43, quinacrine, anisomycin, mitoxantrone, digitoxin, and pyrvinium – strongly reduced SG formation in all three stress contexts (Figures 4A-C). Three of these compounds – quinacrine, mitoxantrone, and pyrvinium – are molecules with planar moieties. The remaining four compounds – 8-hydroxyquinoline, daunorubicin, penfluridol, and benzethonium – strongly reduced SG formation in two of three stress contexts (Figures 4A-C). We concluded that our hit compounds are able to recapitulate their inhibition of SGs in patient-derived iPS-MNs.

**Figure 4.**
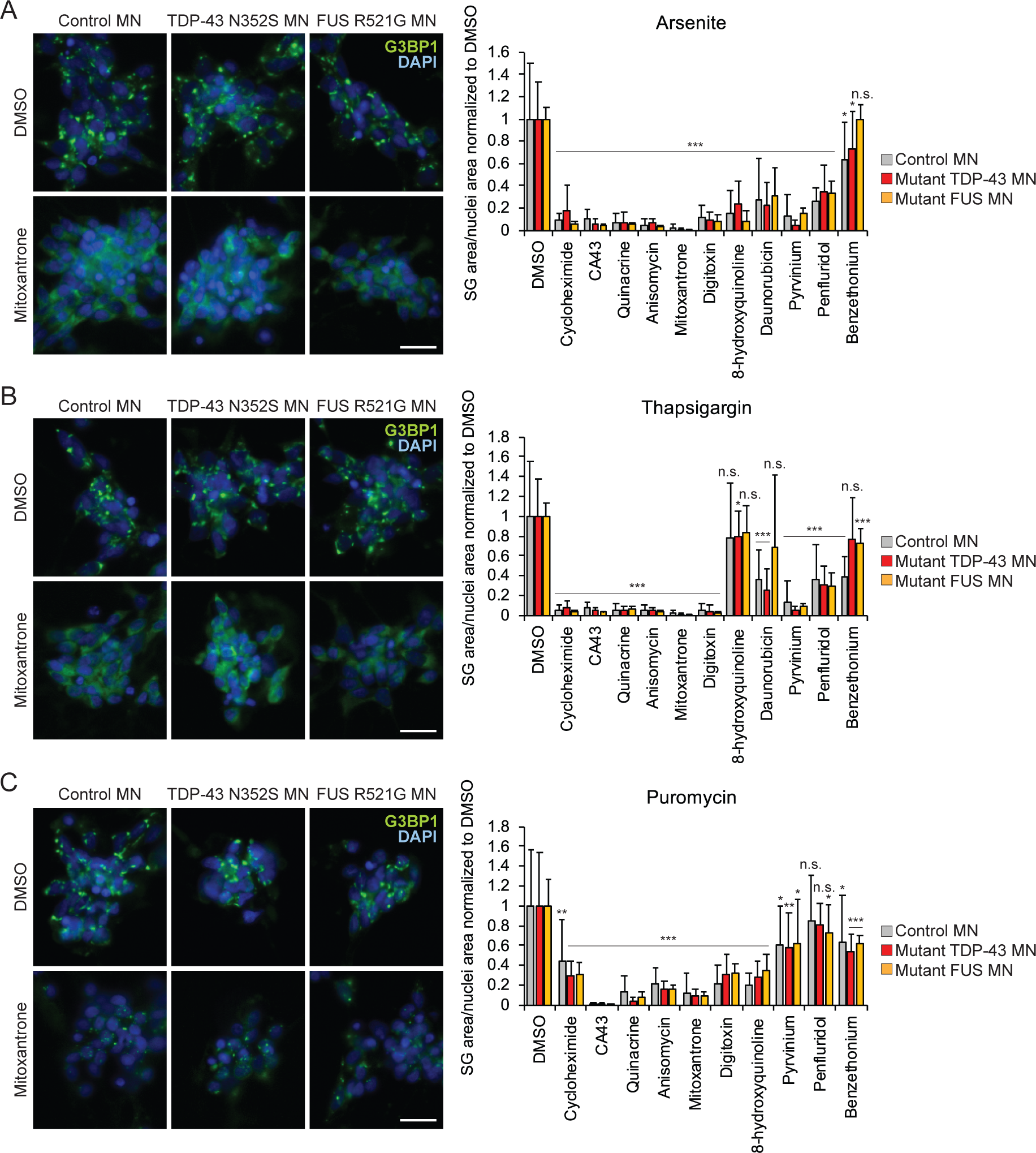
Screen hit compounds inhibit SG formation in iPSC-derived MNs. (A) Representative wide-field fluorescence microscopy of control iPS-MN line and iPS-MNs lines carrying ALS-associated mutations in *TARDBP* and *FUS*, respectively, treated with a nucleic acid intercalating molecule mitoxantrone versus DMSO control under NaAsO_2_ stress. Histograms showing the reduction in SG area/nuclei area in four control lines, four *TARDBP* mutant lines, and two *FUS* mutant lines treated with hit compounds under NaAsO_2_ stress. MN: motor neuron, SG: stress granule. (B) Representative wide-field fluorescence microscopy of control iPS-MN line and iPS-MNs lines carrying ALS-associated mutations in *TARDBP* and *FUS*, respectively, treated with a nucleic acid intercalating molecule mitoxantrone versus DMSO control under thapsigargin stress. Histograms showing the reduction in SG area/nuclei area in four control lines, four *TARDBP* mutant lines, and two *FUS* mutant lines treated with hit compounds under thapsigargin stress. (C) Representative wide-field fluorescence microscopy of control iPS-MN line and iPS-MNs lines carrying ALS-associated mutations in *TARDBP* and *FUS*, respectively, treated with a nucleic acid intercalating molecule mitoxantrone versus DMSO control under puromycin stress. Histograms showing the reduction in SG area/nuclei area in four control lines, four *TARDBP* mutant lines, and two *FUS* mutant lines treated with hit compounds under puromycin stress. (A-C) Histogram values are normalized to cells treated with the vehicle control DMSO. Significance levels are from two-tailed two sample Student’s t-tests to DMSO control: * p < 0.05, ** p < 0.01, *** p < 0.001. Error bars are sample standard deviations from at least five biological replicates of four control lines, four *TARDBP* mutant lines, and two *FUS* mutant lines. Scale bars are 20 μm.

### ALS-associated RNA-binding Proteins are Recruited as SG Shells

Contemporaneous with our small molecule screen, we utilized our G3BP1-GFP reporter lines to evaluate if ALS-associated RBPs such as TDP-43, FUS, HNRNPA2B1 and TIA1 were present in SGs and if so, in which subcompartment i.e. SG core or shell. Using a previously reported protocol (*10*), we isolated SG-enriched fractions from our G3BP1-GFP expressing HEK293xT cells and NPCs. As expected, we found that fractions isolated from NaAsO_2_-stressed cells contained distinct GFP-positive spherical bodies, whereas those from unstressed cells showed diffuse GFP signal (Figure S5A). Additionally, gel electrophoresis and subsequent silver staining of these fractions from stressed and unstressed cells revealed changes in band intensity for numerous proteins upon NaAsO_2_ stress, likely representing increased levels of SG shell proteins and proportionally decreased levels of SG core proteins as SG composition becomes more complex (Figure S5B).

Next, we performed Western blot analysis of fractions from stressed and unstressed cells to determine the relative abundances of well-known SG-associated proteins, including G3BP1, ATAXIN2, CAPRIN1, UBAP2L, PABPC1, DDX3, and USP10, as well as ALS-associated RBPs, including TDP-43, FUS, HNRNPA2B1, and TIA1. Surprisingly, we found that the relative abundances of SG-associated proteins in these fractions decreased upon NaAsO_2_ stress, whereas the relative abundances of the four ALS-associated proteins TDP-43, FUS, HNRNPA2B1, and TIA1 increased upon NaAsO_2_ stress (Figures 5A-B). CAPRIN1, UBAP2L, and USP10 are SG proteins that have been shown to directly interact with G3BP1 and can affect SG formation (*11, 12, 36*). ATAXIN2 binds mRNAs and altered levels of this protein can disrupt SG formation, while PABPC1 and DDX3 are ubiquitous RBPs that are bound to the 3’UTR and 5’UTR of most mRNAs, respectively (*46, 49*). Supporting and further extending recent proximity-labeling proteomic studies of SGs (*11, 12*), our Western blot results indicate that SG-associated proteins already pre-exist in a core-like complex with G3BP1 even in the absence of stress but other proteins such as ALS-associated RBPs are further recruited to SGs (and are present in the shell) during stress.

**Figure 5.**
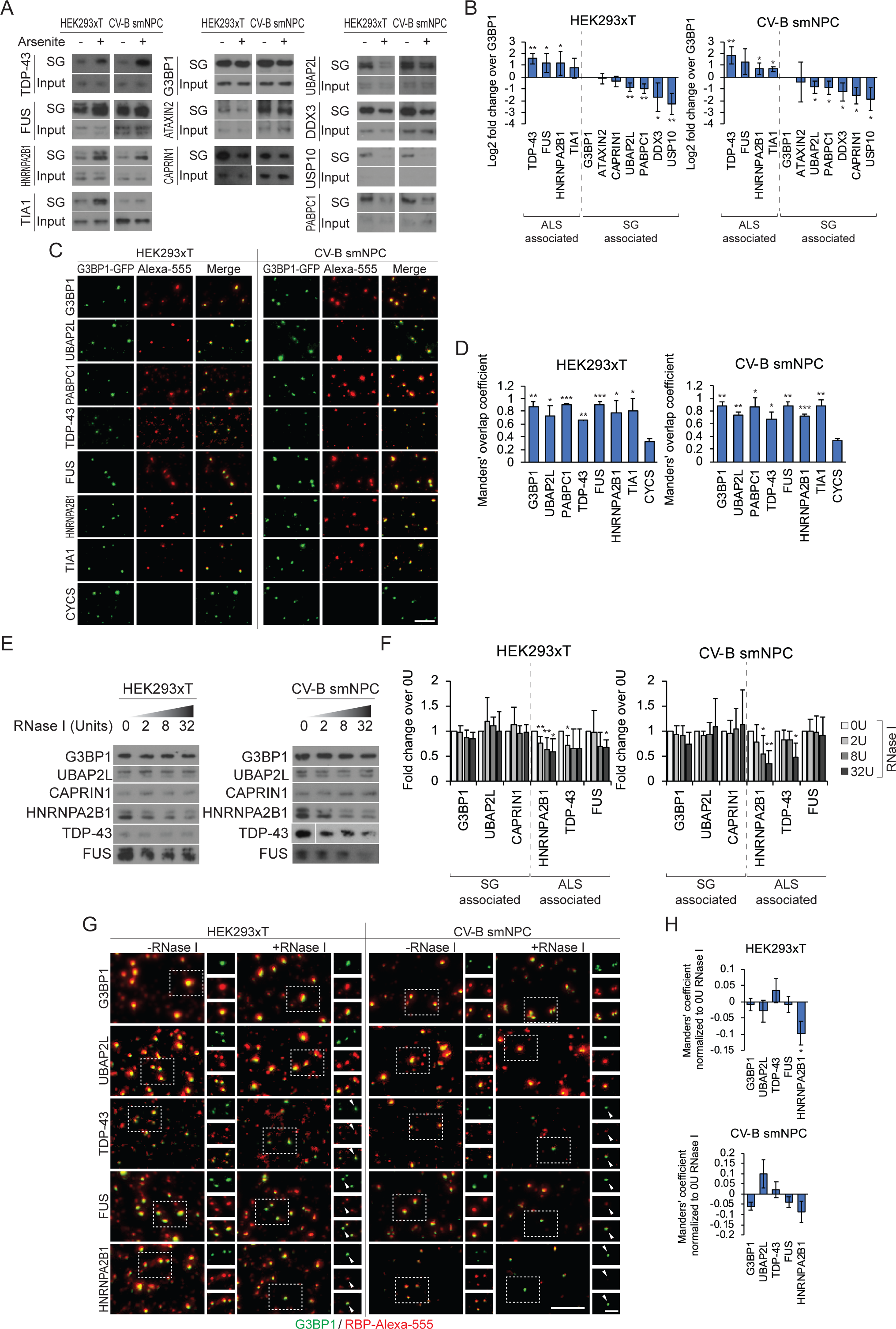
ALS-associated RBPs assemble onto SGs in part through RNA-RBP interactions. (A) Representative Western blots of ALS-associated RBPs and SG-associated proteins in SG-enriched fractions or whole-cell lysate input from unstressed versus NaAsO_2_ -stressed cells. SG-enriched fractions were isolated from G3BP1-GFP HEK293xT cells and CV-B small molecule neural precursor cells. SG: stress granule, NPC: small molecule neural precursor cells. (B) Histograms quantifying the Western blotting in (A) of ALS-associated RBPs and SG-associated proteins in SG-enriched fractions from unstressed versus NaAsO_2_ -stressed cells. Histogram values are ratios of Western blot band intensities from stressed fractions divided by unstressed fractions, normalized to G3BP1, and log_2_-transformed. Significance levels are from two-tailed one sample Student’s t-tests: * p < 0.05, ** p < 0.01. Error bars are sample standard deviations from three experimental replicates. (C) Representative wide-field fluorescence microscopy of NaAsO_2_ SG-enriched fractions probed by immunofluorescence for ALS-associated RBPs and SG-associated proteins. G3BP1 was probed as a positive control and CYCS was probed as a negative control. Scale bar is 10 μm. (D) Histograms quantifying the co-localization in (C) of G3BP1-GFP SGs and immunofluorescence probing of ALS-associated RBPs and SG-associated proteins in NaAsO_2_ SG-enriched fractions. Significance levels are from two-tailed two sample Student’s t-tests to CYC: * p < 0.05, ** p < 0.01, *** p < 0.001. Error bars are sample standard deviations from three experimental replicates. (E) Representative Western blots of ALS-associated RBPs and SG-associated proteins in NaAsO_2_ SG-enriched fractions after digestion with increasing amounts of RNase I. (F) Histograms quantifying the Western blotting in (E) of ALS-associated RBPs and SG-associated proteins in NaAsO_2_ SG-enriched fractions after digestion with increasing amounts of RNase I. Histogram values are normalized to samples treated with 0 Units of RNase I. Significance levels are from two-tailed one sample Student’s t-tests: * p < 0.05, ** p < 0.01. Error bars are sample standard deviations from three experimental replicates. (G) Representative wide-field fluorescence microscopy of NaAsO_2_ SG-enriched fractions probed by immunofluorescence for ALS-associated RBPs and SG-associated proteins, in the absence or presence of RNase I digestion. Scale bar is 10 μm (5 μm for inset). (H) Histograms quantifying the co-localization in (G) of G3BP1-GFP SGs and immunofluorescence probing of ALS-associated RBPs and SG-associated proteins in NaAsO_2_SG-enriched fractions, in the absence or presence of RNase I digestion. Histogram values are normalized to samples treated with 0 Units of RNase I. Significance levels are from two-tailed one sample Student’s t-tests: * p < 0.05. Error bars are estimated standard error of the means of 18 biological replicates from two experimental replicates.

Immunofluorescence probing of SG-enriched fractions from NaAsO_2_ -stressed HEK293xT cells and NPCs also confirm that ALS-associated RBPs join G3BP1 subcomplexes under stress. We found significant co-localization between G3BP1-GFP-positive SGs and ALS-associated proteins TDP-43, FUS, HNRNPA2B1, and TIA1 (Figures 5C-D). As positive controls, we found significant co-localization for several SG-associated proteins including UBAP2L, PABPC1, CAPRIN1, and USP10 (Figures 5C-D and S5C). The negative control CYCS did not co-localize with SGs in these fractions (Figures 5C-D). These results confirm that ALS-associated RBPs such as TDP-43, FUS, HNRNPA2B1, and TIA1 assemble onto G3BP1-positive SG cores.

### Assembly of ALS-associated RBPs onto SGs is RNA-dependent

Since SGs contain RNA and the ALS-associated proteins TDP-43, FUS, and HNRNPA2B1 are RBPs, we hypothesized that assembly of these proteins is RNA-dependent. Indeed, we showed by Western blotting that RNase I digestion of fractionated SG-enriched fractions from NaAsO_2_ -stressed HEK293xT cells and NPCs led to reduction of HNRNPA2B1 from SGs in a dose-dependent manner in both cell types. We also found that RNase I digestion reduced TDP-43 and FUS from SGs in fractions from NPCs and HEK293xT cells, respectively (Figures 5E-F). This is in contrast to the SG-associated RBPs UBAP2L and CAPRIN1, which were resistant to RNase I digestion (Figures 5E-F).

Immunofluorescence staining of SG-enriched fractions in the absence or the presence of RNase I digestion indicated that co-localization of HNRNPA2B1 with G3BP1-GFP-positive SGs decreased following RNase I digestion, but not co-localization of G3BP1 or UBAP2L (Figures 5G-H). Although we were not able to detect decreases in co-localization of TDP-43 or FUS with G3BP1-GFP-positive SGs following RNase I digestion, by first incubating SG-enriched fractions with 1,6-hexanediol to disrupt hydrophobic interactions followed by RNase I digestion, we demonstrate decreased co-localization of FUS with G3BP1-GFP-positive SGs (Figures S5D). These Western blot and immunofluorescence microscopy results suggest that the assembly of HNRNPA2B1, and to a lesser extent TDP-43 and FUS, onto SGs is RNA-dependent.

### Molecules with Planar Moieties Reduce ALS-associated RBPs from SGs

As our small molecule screen demonstrated that molecules with planar moieties, such as mitoxantrone, doxorubicin, and daunorubicin, decreased the average size of SGs while increasing the number of SGs per cell (Figure S2A), we reasoned that these molecules likely modulate SG fusion by interfering with proper formation of SG shells (*9*). As these planar moiety-containing molecules had been reported to be nucleic acid intercalating molecules that directly bind to RNA (*50–55*), we further reasoned that these compounds may reduce the RNA-dependent assembly of ALS-associated RBPs into SGs.

Indeed, immunofluorescence probing of SG-enriched fractions incubated in the absence or the presence of mitoxantrone demonstrated a significant decrease of co-localization of TDP-43, FUS, and HNRNPA2B1 with G3BP1-GFP-positive SGs. Co-localization of G3BP1 or UBAP2L was not affected (Figures 6A-B). Similarly, we observed a reduction of co-localization of FUS and HNRNPA2B1 with G3BP1-GFP-positive SGs following incubation of SG-enriched fractions with doxorubicin (Figure S6A). In contrast, incubation of SG-enriched fractions with another compound, CA43, which does not have planar moieties, did not result in decreased co-localization of TDP-43, FUS, and HNRNPA2B1 with G3BP1-GFP-positive SGs (Figure S6B).

**Figure 6.**
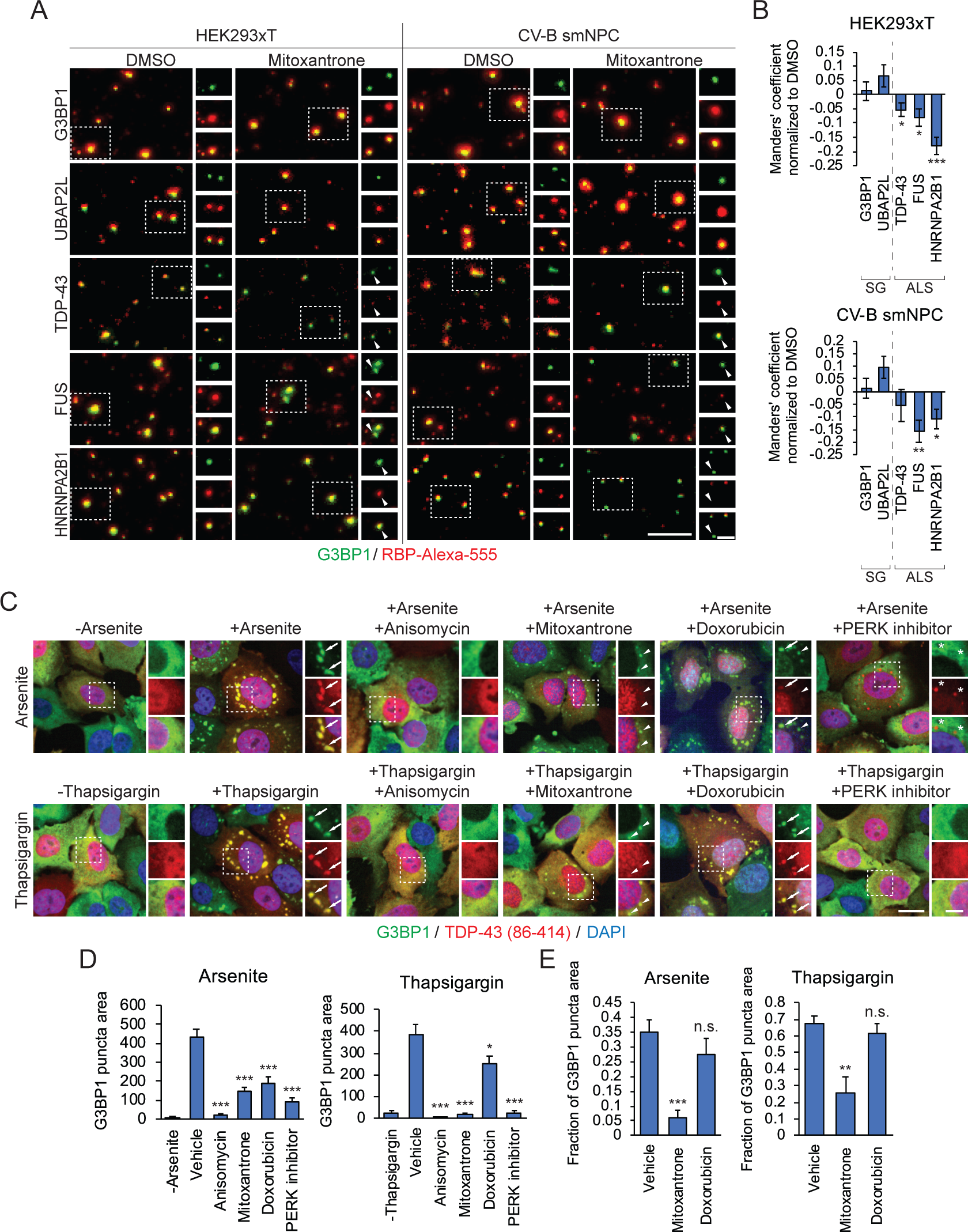
Assembly of ALS-associated RBPs can be disrupted by molecules with planar moieties. (A) Representative wide-field fluorescence microscopy of NaAsO_2_ SG-enriched fractions probed by immunofluorescence for TDP-43 in the absence or presence of overnight incubation with 100 μM mitoxantrone. Scale bar is 10 μm (5 μm for inset). NPC: small molecule neural precursor cells, RBP: RNA-binding protein. (B) Histograms quantifying the change in co-localization in (A) of G3BP1-GFP SGs and immunofluorescence probing of TDP-43 in NaAsO_2_ SG-enriched fractions, in the absence or presence of overnight incubation with 100 μM mitoxantrone. Histogram values are normalized to samples treated with the vehicle control DMSO. Significance levels are from two-tailed one sample Student’s t-tests: * p < 0.05, ** p < 0.01, *** p < 0.001. Error bars are estimated standard error of the means from 18 biological replicates from two experimental replicates. SG: stress granule. (C) Representative wide-field fluorescence microscopy of H4 cells transiently expressing G3BP1-mCherry (false-colored green for consistency with other fluorescent imaging of G3BP1) and GFP-TDP-43ΔNLS (amino acid residues 86-414/no nuclear localization signal; false-colored red), stressed with NaAsO_2_ or thapsigargin, in the absence and presence of SG-modulating compounds. Arrows denote co-localized G3BP1-mCherry SG-like puncta and GFP-TDP-43ΔNLS foci. Arrowheads denote G3BP1-mCherry puncta that are largely devoid of co-localized GFP-TDP-43ΔNLS. Stars denote GFP-TDP-43ΔNLS foci that are largely devoid of co-localized G3BP1-mCherry. Scale bar is 50 μm (20 μm for insets). (E) Histograms quantifying the formation in (C) of G3BP1-mCherry SG-like puncta in H4 cells transiently expressing G3BP1-mCherry and GFP-TDP-43ΔNLS and stressed with NaAsO_2_ or thapsigargin, in the absence or presence of SG-modulating compounds. Significance levels are from two-tailed two sample Student’s t-tests to vehicle control: * p < 0.05, *** p < 0.001. Error bars are estimated standard error of the means from n ≥ 35 cells. (E) Histograms quantifying in (C) the fraction of G3BP1-mCherry SG-like puncta that have GFP-TDP-43ΔNLS co-localization in H4 cells transiently expressing G3BP1-mCherry and GFP-TDP-43ΔNLS and stressed with NaAsO_2_ or thapsigargin, in the absence or presence of SG-modulating compounds. Significance levels are from two-tailed two sample Student’s t-tests to vehicle control: ** p < 0.01, *** p < 0.001. Error bars are estimated standard error of the means from n ≥ 14 cells.

Western blotting analysis of SG-enriched fractions incubated with mitoxantrone and daunorubicin also indicated reduction of TDP-43 from SGs (Figure S6D). Two other compounds with planar moieties, pyrvinium and pararosaniline, additionally reduced FUS and HNRNPA2B1 from SGs in fractions from HEK293xT cells, and reduced TDP-43 from SGs in fractions from NPCs (Figure S6C). By contrast, SG proteins G3BP1 and CAPRIN1 were not reduced from SGs during overnight incubation with any of these SG-modulating compounds with planar moieties (Figure S6C). Incidentally, many of these compounds appear brightly colored under visible light due to the presence of aromatic rings with extended conjugated pi-bond systems, and we were readily able to observe their association with pelleted SGs with the naked eye (Figure S6D). In summary, these results indicate that molecules with planar moieties, such as the nucleic acid intercalators mitoxantrone, daunorubicin, pyrvinium, and pararosaniline, reduce ALS-associated RBPs from SGs *in vitro*.

### Planar Moiety-containing Compounds Inhibit TDP-43 Accumulation in SGs in TDP-43ΔNLS cells

Since compounds with planar moieties reduce TDP-43 from SGs *in vitro*, we next evaluated these compounds in disease-relevant cellular models that harbor mislocalized TDP-43 protein. In H4 neuroglioma cells that express G3BP1-mCherry and GFP-TDP-43ΔNLS (86-414), NaAsO_2_ or thapsigargin treatment strongly induced formation of G3BP1-mCherry SGs as well as GFP-TDP-43ΔNLS foci that essentially co-localized completely with the G3BP1-mCherry SGs (Figures 6C-D and S6E).

As expected from our results above, pre-treatment with each of four SG inhibiting compounds anisomycin, mitoxantrone, doxorubicin, or GSK2606414 significantly reduced the amount of SG formation per cell (Figures 6C-D). Excitingly, pre-treatment with mitoxantrone and doxorubicin, which have planar moieties strongly inhibited co-localization of GFP-TDP-43ΔNLS with residual SGs (Figures 6C and 6E). In contrast, pre-treatment with PERK inhibitor GSK2606414 inhibited SG formation but had no effect on cytoplasmic GFP-TDP-43ΔNLS foci, which were not co-localized with G3BP1-mCherry (Figures 6C-D and S6E). These results demonstrate that planar nucleic acid intercalators inhibit accumulation of TDP-43 in SGs in live cells.

### Planar Moiety-containing Compound Reduces Puromycin-induced Persistent Cytoplasmic TDP-43

Next, we evaluated if molecules with planar moieties mitigate ALS-associated phenotypes using iPS-MNs. First, we determined whether SG formation leads to persistent cytoplasmic accumulation of TDP-43 in motor neurons, as has been reported in HeLa cells (*56–60*). We treated *TARDBP* mutant and control iPS-MNs with puromycin and then washed out the puromycin to allow the cells to recover from stress. We found that during stress, significantly more TDP-43 accumulated in SGs in the *TARDBP* mutant iPS-MNs than in the controls (Figures 7A-B). During recovery from stress following washout of puromycin, significantly more TDP-43 remained localized in the cytoplasm of the *TARDBP* mutants than in the controls (Figures 7A-B). This cytoplasmic TDP-43 persisted for at least 24 hours after washout of the stressor. Furthermore, biochemically isolated SG-enriched fractions from mutant IPS-MNs contained significantly more TDP-43, FUS, and G3BP1 than fractions from control IPS-MNs during recovery after stress (Figures 7C and S7A-D). Similarly, significantly more TDP-43 accumulated in SGs in IPS-MNs harboring ALS-associated *FUS* mutations than in the control IPS-MNs (Figures 7D-E). During recovery from stress, we observed a dramatic increase in cytoplasmic TDP-43 in the *FUS* mutant IPS-MNs compared to controls (Figures 7D-E). This increase in cytoplasmic TDP-43 in the *FUS* mutant IPS-MNs manifested in roughly 10-20% of cells as large cytoplasmic puncta that did not co-localize with G3BP1 (Figure 7D).

**Figure 7.**
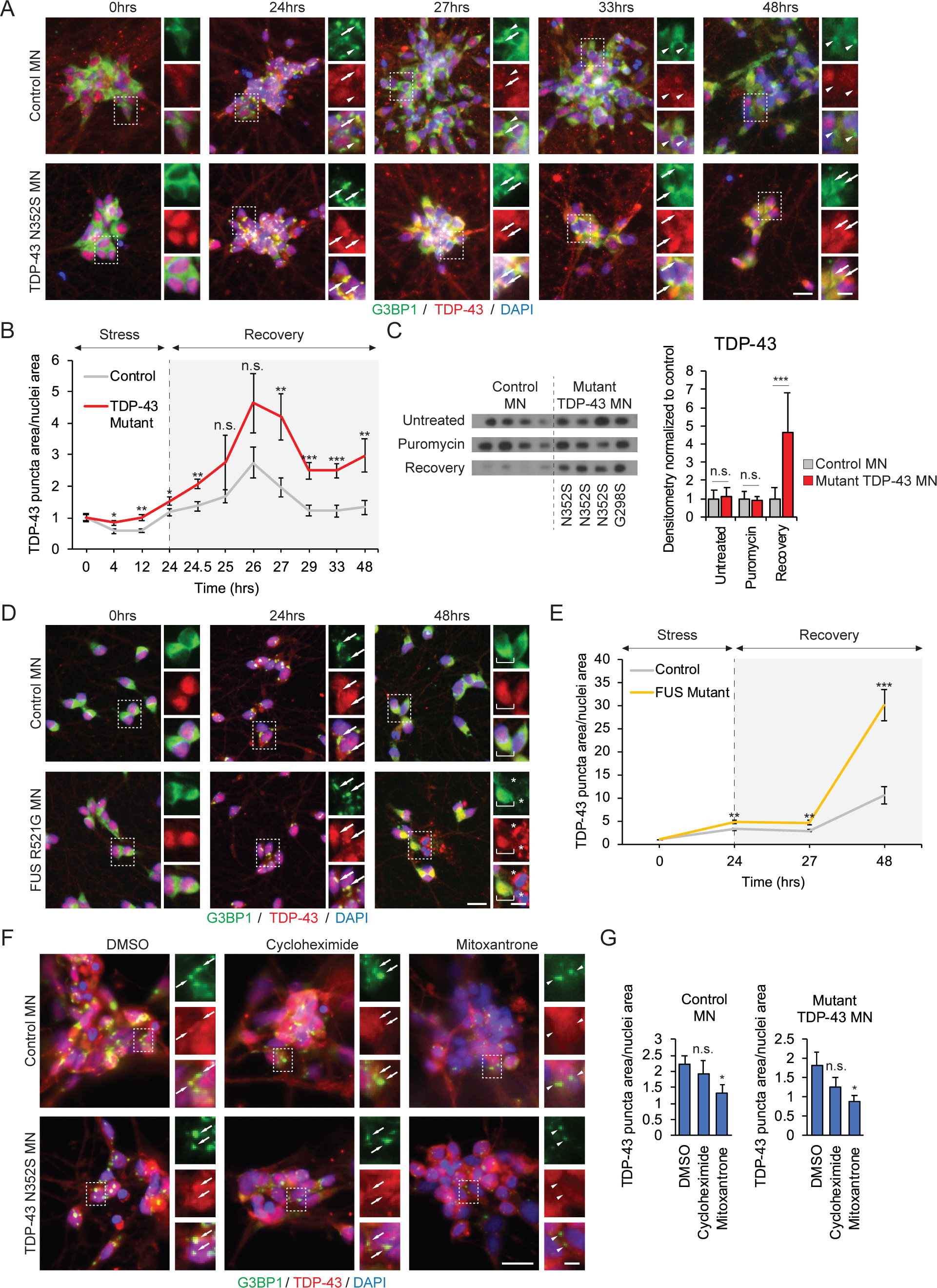
Mutant MNs exhibit persistent cytoplasmic TDP-43 following puromycin stress and washout. (A) Representative wide-field fluorescence microscopy of iPS-MNs probed by immunofluorescence for G3BP1 and TDP-43 and stained with DAPI at several time points during puromycin stress and puromycin washout. Arrows denote co-localized G3BP1 SGs and TDP-43 puncta. Arrowheads denote G3BP1 SGs that are largely devoid of co-localized TDP-43. Scale bar is 20 μm (10 μm for insets). MN: motor neuron. (B) Scatterplot quantifying the formation in (A) of cytoplasmic TDP-43 foci in iPS-MNs from individuals carrying ALS-associated mutations in *TARDBP* or control MNs. Scatterplot values are normalized to samples at the 0 hour time point. Significance levels are from two-tailed two sample Student’s t-tests to control MNs: * p < 0.05, ** p < 0.01, *** p < 0.001. Error bars are estimated standard error of the means from five biological replicates of four control lines and four *TARDBP* mutant lines. (C) Representative Western blots and histogram quantifying the amount of TDP-43 in SG-enriched fractions isolated from *TARDBP* mutant or control iPS-MNs under no stress conditions, after 24 hours of puromycin stress, or after 24 hours of washout following 24 hours of puromycin stress. Histogram values are normalized to the control line samples. Significance levels are from two-tailed two sample Student’s t-tests to control MNs: * p < 0.05. Error bars are sample standard deviations of three biological replicates across four control lines and four *TARDBP* mutant lines. (D) Representative wide-field fluorescence microscopy of iPS-MNs probed by immunofluorescence for G3BP1 and TDP-43 and stained with DAPI at several time points during puromycin stress and puromycin washout. Arrows denote co-localized G3BP1 SGs and TDP-43 puncta. Brackets denote broad regions of cytoplasmic TDP-43. Stars denote cytoplasmic TDP-43 puncta that are largely devoid of co-localized G3BP1. Scale bar is 20 μm (10 μm for insets). (E) Scatterplot quantifying the formation in (D) of cytoplasmic TDP-43 foci in iPS-MNs from individuals carrying ALS-associated mutations in *FUS* or control MNs. Scatterplot values are normalized to samples at the 0 hour time point. Significance levels are from two-tailed two sample Student’s t-tests to control MNs: ** p < 0.01, *** p < 0.001. Error bars are estimated standard error of the means from five biological replicates of four control lines and two *FUS* mutant lines. (F) Representative wide-field fluorescence microscopy of iPS-MNs probed by immunofluorescence for G3BP1 and TDP-43 and stained with DAPI; iPS-MNs were incubated with SG-inhibiting compounds cycloheximide or mitoxantrone or DMSO control and stressed with puromycin. Arrows denote co-localized G3BP1 SGs and TDP-43 puncta. Arrowheads denote G3BP1 SGs that are largely devoid of co-localized TDP-43. Scale bar is 20 μm (5 μm for insets). (G) Histograms quantifying the formation in (F) of cytoplasmic TDP-43 foci in iPS-MNs from individuals carrying ALS-associated mutations in *TARDBP*, or control iPS-MNs; cells were incubated with SG-inhibiting compounds cycloheximide or mitoxantrone or DMSO control and stressed with puromycin. Significance levels are from two-tailed two sample Student’s t-tests to DMSO control: * p < 0.05. Error bars are estimated standard error of the means from five biological replicates of four control lines and four *TARDBP* mutant lines.

Encouraged that IPS-MNs carrying ALS-associated mutations exhibit a clear ALS-associated molecular phenotype, we next incubated IPS-MNs from controls, *TARDBP* and *FUS* mutants with DMSO control, cycloheximide, or planar compound mitoxantrone in the presence of puromycin stress. As mentioned above, cycloheximide and mitoxantrone strongly inhibited SG formation compared to the DMSO treatment (Figure 4C). Importantly, mitoxantrone also significantly abrogated cytoplasmic localization of TDP-43 in both control and *TARDBP* mutant IPS-MNs, whereas DMSO and cycloheximide largely did not (Figures 7F-G and S7E-F). These results indicate that molecules with planar moieties can prevent the ALS-associated phenotype of persistent cytoplasmic TDP-43 induced by cellular stress.

## DISCUSSION

In this study, a high-content small molecule screen was performed in which we identified approximately 100 compounds consisting of eight compound classes that modulate SG formation, six of which have not previously been reported to affect SGs. These include 50 compounds that act at cell surface targets such as ion channels, receptors, or lipid membranes; seven compounds that modulate inflammatory signaling pathways; and 20 compounds that are molecules with planar moieties. This diversity of SG-modulating compounds greatly expands the toolbox of molecular compounds with which to probe the relationship between SGs and disease. Previous studies of SGs in disease models have relied on a handful of SG modulators: cycloheximide, the PERK inhibitor GSK2606414, or ISRIB, a small molecule SG inhibitor identified in a translation-based screen (*5, 11, 34, 60, 61*). This study offers a host of alternative compounds that act through orthogonal mechanisms which may prove useful in dissecting the role of SGs in disease models.

This study is also the first high-content screen in which compounds (two-thirds of the hits) were identified to not only modulate the overall amount of SG formation, but also SG size and the numbers of SGs per cell. Surprisingly, we observed that molecules with planar moieties such as nucleic acid intercalators not only inhibit SG formation, but also alter SG size and the numbers of SGs per cell in proliferative cells as well as patient-specific and control post-mitotic iPS-MNs. We reasoned that the nuanced effects of these compounds on SGs may reveal the molecular rules underlying SG formation. Specifically, molecules with planar moieties likely act by inhibiting SG shell formation which would lead to defects in SG fusion and maturation across a diverse range of stressors, thus manifesting as smaller and more numerous SGs (*9*). Interestingly, we demonstrate that eight compounds – five of which are molecules with planar moieties – reduce total SG formation without altering phosphorylation of eIF2α. This is surprising since phosphorylation of eIF2α is a committing step towards SG formation. This study therefore also offers SG-inhibiting compounds that act through orthogonal mechanisms, which will be useful in future dissection of the role of SGs in disease models.

To evaluate a potential molecular mechanism for how particular compounds affect specific subcompartments of SGs such as SG shells, we leveraged a recently published biochemical approach (*10*). We fractionated stressed cells and used the SG-enriched fractions as an *in vitro* model system (*10*). Using a combination of silver staining and Western blot analysis, we found that fractions from stressed cells were depleted or enriched for particular proteins compared to fractions from unstressed cells. Among the depleted proteins were proteins reported to be SG-associated proteins such as CAPRIN1, UBAP2L, and USP10, whereas the enriched proteins included ALS-associated proteins such as TDP-43, FUS/TLS, HNRNPA2B1 and TIA1. We observed that the depleted proteins consist of components of cores in SG that is pre-existing in mRNP complexes prior to cellular stress (*11*). The relative representation of these SG core proteins is reduced upon stress due to assembly of a complex array of SG shell proteins onto the cores. These SG shell proteins are enriched for the ALS-associated proteins whose representation becomes higher relative to the SG core proteins. This represents the first look into differences in protein compositions between cores and shells, and paves the way for systematic enumeration of core versus shell proteins as well as functional high-content screens to assess the differential contributions of core and shell proteins to SG formation.

By treating SG-enriched fractions with RNase I, we further demonstrated that the assembly of several of these SG shell proteins is RNA-dependent, revealing for the first time how SG shell proteins may be recruited to SGs. This is consistent with recent RNA-seq analyses of SG-enriched fractions where RNAs in these preparations were enriched for the RNA binding motifs of putative SG shell proteins TDP-43 and TIA1 (*3*). Furthermore, a recent systematic survey of the protein interactome for HNRNPA2B1 revealed that this RBP associates with a large number of other RBPs indirectly via RNA-RBP interactions (*13*). Similarly, HNRNPA2B1 may indirectly associate with SG core proteins via RNA-RBP interactions. Finally, it has been recently reported that RNA can regulate the phase separation of RBPs, consistent with the hypothesis that SGs may be assembled in part through phase separation processes (*62, 63*).

Finally, comparing incubation of SG-enriched fractions with mitoxantrone, doxorubicin, or CA43 to incubation with DMSO control, we found that mitoxantrone and doxorubicin, but not CA43 or DMSO, dislodged the putative SG shell proteins TDP-43, FUS, and HNRNPA2B1 from SGs. Given that mitoxantrone and doxorubicin are both nucleic acid intercalating molecules reported to directly interact with RNA, we reasoned that these two compounds interfere with RNA-RBP interactions to dislodge putative SG shell proteins (*47, 50–53*). IDRs and PrLDs are often enriched in amino acids with large aromatic side chains (*64*), and coacervation of charged aromatic side chains help regulate phase separation of IDR-containing proteins. Mitoxantrone, doxorubicin, and other molecules with planar moieties may interact with these aromatic side chains via π-π stacks (*2, 65*). For example, acridine derivatives such as quinacrine are largely planar molecules that directly interact with prion protein and amyloid beta inclusions (*48, 66*). Given that SG formation has been reported to cause altered nucleocytoplasmic shuttling and localization of ALS-associated RBPs (*60*), these compounds which can dislodge ALS-associated RBPs from SGs raises the possibility that these compounds represent a new therapeutic approach for disease.

To investigate this hypothesis, we developed a model system of ALS/FTD, in which iPS-MNs are exposed to 24h of stress by puromycin treatment, followed by washout and recovery in medium without puromycin. We demonstrated increased cytoplasmic accumulation of TDP-43 during puromycin stress, which persisted for at least 24 hours into the recovery period in *TARDBP* and *FUS* mutant, but not in control iPS-MNs. This represents the first disease model showing motor neurons exhibit a persistent molecular TDP-43 phenotype resembling the cytoplasmic mislocalization of TDP-43 in patient tissues (*56–59*), and ties together SGs with disease pathophysiology. Excitingly, when we treated the cells with mitoxantrone, a molecule with planar moieties, we not only reduced SG formation, but also reduced cytoplasmic accumulation of TDP-43. Further corroborating these results, we found that mitoxantrone also decreased association of TDP-43 with SGs in H4 cells engineered with a TDP-43ΔNLS (86-414) expression cassette.

Overall, our results in this study point us towards a two-step model for SG formation (Figure 8): In the first step, cellular stress signaling leads to phosphorylation of eIF2α, freezing ribosomes in the 48S pre-initiation stage at the 5’ untranslated regions (UTRs) of mRNAs. Actively translating polysomes eventually run off the ends of mRNAs, while at the same time the SG scaffold protein G3BP1 can directly bind to the 40S ribosome subunit. G3BP1 can undergo homotypic interactions via IDRs near the C-terminus and catalyze the formation of a SG core. In the second step of SG formation, RBPs assemble onto SGs through RNA-dependent interactions. Many of these RBPs contain IDRs, which can also mediate assembly of additional proteins through heterotypic IDR contacts to form a SG shell in a phase-transition process. We posit that this second step of SG formation is most relevant to disease pathophysiology. Mutations in the IDRs of RBPs or other SG shell proteins may accumulate in local high concentrations in the SG shell, and over a lifetime of stress, SG shells could nucleate the formation of pathological aggregations of mutant proteins, such as those seen in the spinal cord MNs of ALS patients (*29*). In this context molecules with planar moieties such as nucleic acid intercalating molecules directly interact with RNAs, disrupting recruitment of ALS-associated RBPs to the SG shell. Thus, disruption of RNA-RBP interactions in SG shells is a potential therapeutic strategy for ALS/FTD.

**Figure 8.**
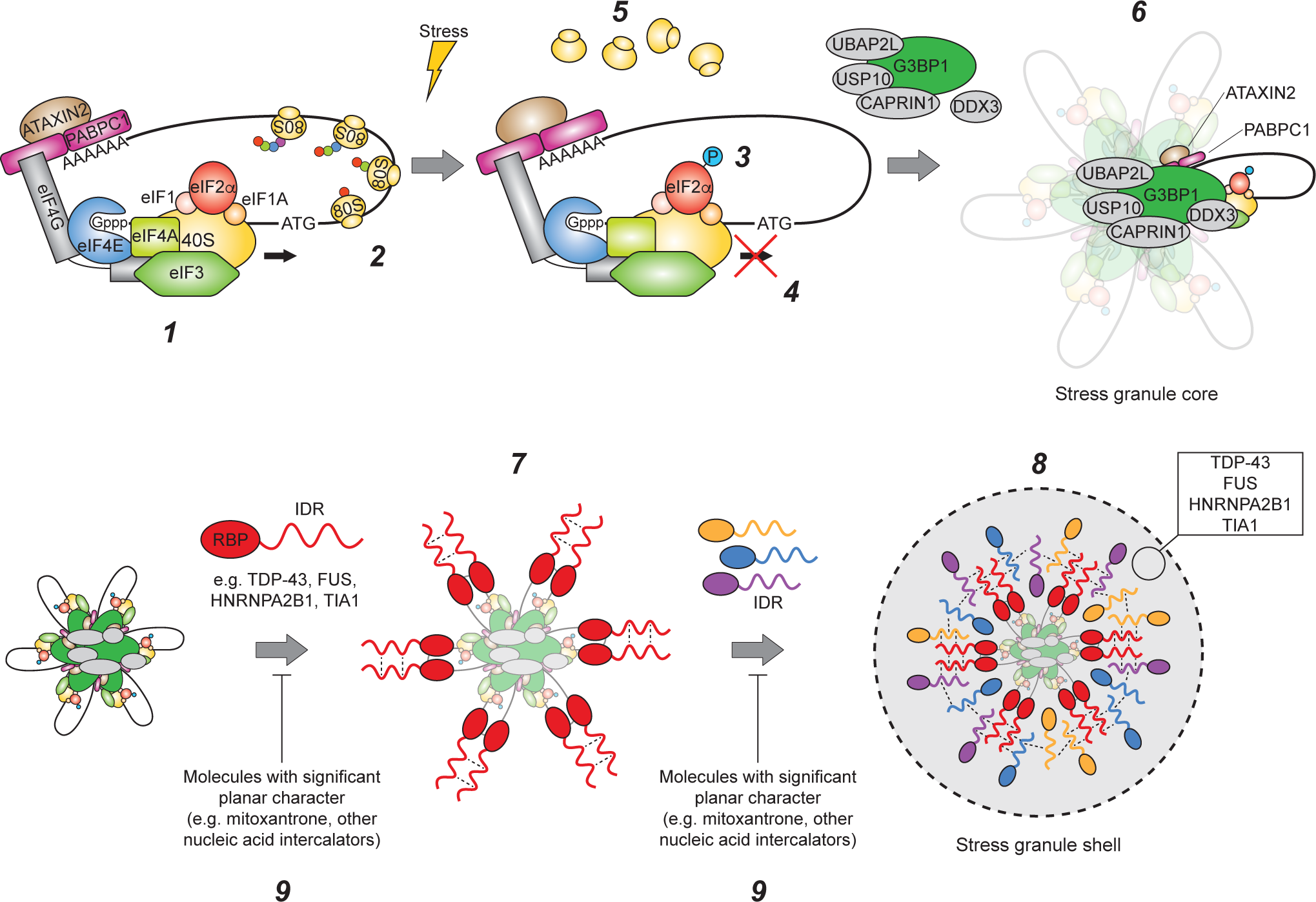
Model relating SG formation and disease pathophysiology. Model relating SG formation and disease pathophysiology Under unstressed conditions, the 48S pre-initiation ribosome complex assembles on the 5’ cap of actively translation mRNAs (1) and scans the 5’ untranslated region for the start codon, upon which the full 80S ribosome assembles and translates the coding sequence (2). Following cellular stress such as exposure to NaAsO_2_ or thapsigargin, stress-activated kinases phosphorylate eIF2α on serine 51 (3), freezing ribosomes at the 48S pre-initiation complex stage at the 5’ cap (4). Elongating 80S ribosomes eventually fall off transcripts (5). Classical SG-associated RBPs such as G3BP1, TIA1, UBAP2L, USP10, CAPRIN1, and DDX3 can bind to the 40S ribosome subunit, the 5’ or 3’ untranslated regions, or G3BP1, and these proteins mediate assembly of several of these ribonucleoproteins into a core SG structure, possibly through a combination of protein-protein interactions via well-ordered protein domains as well as liquid-liquid phase separation interactions of the intrinsically disordered regions of these proteins (6). This core structure contains mRNA whose coding sequences may act as landing sites for the assembly of RBPs, including ALS-associated proteins such as TDP-43, FUS, and HNRNPA2B1 (7). The intrinsically disordered regions (IDR) of these RBPs may then further mediate assembly of other proteins with intrinsically disordered regions, which may or may not be RBPs, forming a shell SG structure (8). This shell structure may lead to a local high concentration of mutant protein in cells carrying mutations in the intrinsically disordered regions of ALS-associated proteins such as TDP-43, and may nucleate the formation of more insoluble and potentially cytotoxic aggregates. (9) SG screen hit compounds disrupt association of ALS-associated proteins with the SG shell, potentially reducing the nucleation and formation of insoluble protein aggregates.

## AUTHOR CONTRIBUTIONS

Conceptualization, M.Y.F., S.M. and G.W.Y; Methodology, M.Y.F., S.M., W.E.D., A.Q.V. and P.J.B.; Formal Analysis, M.Y.F., S.M., W.E.D., P.J.B.; Investigation, M.Y.F., S.M., W.E.D., A.Q.V. and P.J.B.; Writing – Original Draft, M.Y.F.; Writing – Review & Editing, M.Y.F., S.M., W.E.D., J.W.L., and G.W.Y.; Supervision, G.W.Y.; Funding Acquisition, M.M.M., J.W.L., and G.W.Y.

## ACKNOWLEDGEMENTS

We acknowledge members of the G.W.Y. lab for critical comments. We thank Gabriel Pratt and Olga Botvinnik for advice with bioinformatics analyses and Julia Nussbacher and Stefan Aigner for valuable comments on the manuscript. M.Y.F. was supported by an NIH Ruth L. Kirschstein National Research Service Award Institutional Research Training Grant T32 DK 7541-30. S.M. was supported by a postdoctoral fellowship from the Larry L. Hillblom Foundation (2014-A-027-FEL). This work was partially supported by grants from the NIH (NS103172 and HG004659), TargetALS (to G.W.Y. and J.W.L.) and the ALS Association (to G.W.Y.).

## METHODS

### Generating induced pluripotent stem cells from fibroblasts of individuals carrying ALS-associated mutations in *TARDBP* or *FUS*

Human iPSC lines carrying ALS-associated mutations in *TARDBP* (N352S) and control individuals were previously reprogrammed from primary fibroblasts obtained by Dr. John Ravits (University of California, San Diego), as described (*67*). Lines carrying *TARDBP* (G298S) mutations were obtained courtesy of Kevin Eggan, as described (*68*). Lines carrying ALS-associated FUS (R521G) mutations were previously reprogrammed from primary fibroblasts obtained by Franca Cambia, Edward Kasarskis, and Haining Zhu (University of Kentucky), as described (*69*). Informed consent was obtained from all individuals prior to biopsy, and the use of human fibroblasts for this project was approved by the University of California, San Diego Institutional Review Board.

### Generation of human induced pluripotent stem cell-derived small molecule neural precursor cells

Generation of small molecule neural precursor cells (NPC) from human IPSCs was adapted from a previously reported protocol (*70*). Briefly, iPSCs were dissociated into single cells with Accutase and 3×10^6^ cells were seeded into one well of an uncoated 6-well plate. The plate was incubated on an orbital shaker at 90 rpm at 37°C, 5% CO_2_ and the cells were grown in DMEM/F12, GlutaMAX supplement media containing N-2 Supplement (1:100 v/v), CTS B27 Supplement, XenoFree (1:50 v/v), L-ascorbic acid (150 μM), CHIR99021 (3 μM), purmorphamine (0.5 μM), Y-27632 (5 μM), as well as two small molecule SMAD inhibitors: SB431542 (10 μM) and dorsomorphin (1 μM). After 2 days, embryoid bodies (EB) formed, at which point 1/3 of the EBs were transferred to three wells of an uncoated 6-well plate to prevent overgrowth; Y-27632 was also withdrawn from the media. Following this passaging, half media changes were performed every 24 hours. At 6 days, formation of neuroepithelium was evident in the EBs, and a full media change was performed to media that omits SB431542 and dorsomorphin (NPC media, also described below). At 8 days, individual EBs were picked and triturated into small fragments. These EB fragments were seeded onto 10 cm plates coated with Matrigel. These plates were prepared by adding 6 mL of Matrigel (diluted 1:60 v/v in DMEM/F12, GlutaMAX supplement) to each 10 cm plate and incubating for 60 min. EB fragments were maintained on these Matrigel plates as adherent cultures. After 4 days, these adherent EB fragments were dissociated into single cells with Accutase and passaged at a ratio of 1:6 to 1:8 onto new Matrigel plates. Every 3-6 days thereafter these cultures were passaged at a 1:10 to 1:15 ratio. After 5-6 passages, most non-NPCs disappear and homogeneous colonies of NPCs remained. These NPCs may be passaged at least 20 times at a 1:5 to 1:15 ratio.

### Generation of human iPS-MNs

Generation of iPS-MNs was adapted from a previous reported protocol (*23*). Briefly, iPSCs were dissociated into single cells with Accutase and 1×10^6^ cells were seeded into one well of a Matrigel-coated 6-well plate. These Matrigel-coated plates were prepared by adding 1 mL of Matrigel (diluted 1:60 v/v in DMEM/F12, GlutaMAX supplement) to each well of a 6-well plate and incubating the plate for 60 min. The dissociated iPSCs were grown for 6 days in DMEM/F12, GlutaMAX supplement media containing N-2 Supplement (1:100 v/v), CTS B27 Supplement, XenoFree (1:50 v/v), L-ascorbic acid (150 μM), CHIR99021 (3 μM), as well as two small molecule SMAD inhibitors: SB431542 (10 μM) and dorsomorphin (1 μM). ROCK inhibitor (Y-27632) may also be added to the media to a final concentration of 5 μM to reduce cell death. Full media changes were performed every 24 hours. At 7 days, a full media change was performed in which CHIR99021 was withdrawn from the media and retinoic acid (1.5 μM) and Smoothened Agonist (200 nM) was added to the media. The cells were cultured for a further 8 days in this media with daily full media changes. At 15 days, the cells were dissociated: Cells were washed once with PBS and 3 mL of Accutase was added per well of a 6-well plate. Cells were incubated with Accutase for 30 min. Next, each well of cells was triturated gently 4-5 times, transferred to 15 mL centrifuge tubes containing 6 mL of DMEM/F12, GlutaMAX supplement, and the cells pelleted by centrifuging for 5 min at 1000 g. The cell pellets should be resuspended and 1×10^7^ cells seeded onto each 10 cm Matrigel-coated plate; the media should also be supplemented with 10 μM Y-27632 to reduce cell death. Full media changes were performed every 24 hours. At 20 days, the media was modified to reduce Y-27632 (2 μM) and to include three growth factors: recombinant human BDNF (2 ng/ml), recombinant human CNTF (2 ng/mL), and recombinant human GDNF (2 ng/ml). At 22 days, retinoic acid and Smoothened Agonist were withdrawn from the media and DAPT (2 μM) was added to aid maturation to MNs. At this point, full media changes were performed every 2 days. At 26 days, DAPT was withdrawn from the media. At 28 days, iPS-MN cultures were obtained that stained positive for known MN markers Hb9, Islet-1/2 and SMI-31.

### Plasmid construction and G3BP1-GFP reporter line generation

To generate G3BP1-GFP SG reporter lines, the HR120PA-1 targeting vector (System Biosciences) was modified to include a 1.5 kilobase 5’ arm of homology span part of intron 11 and exon 12 of the endogenous human G3BP1 locus, and a 1.5 kilobase 3’ arm of homology from the 3’ untranslated region of G3BP1. These arms of homology were amplified from genomic DNA and the modified G3BP1 targeting vector was assembled via Gibson assembly. HEK293xT cells were transfected with this targeting vector and the Cas9 expression vector PX458 using lipofectamine 2000. Human induced pluripotent stem cells were electroporated using an Amaxa Nucleofector with Stem Cell Kit 1 and pulse setting B-016. Cells were selected beginning two days after transfection with puromycin at 1 μg/ml for four days. After two weeks, G3BP1-GFP colonies were picked using a steromicroscope.

### Cell culture conditions

Human induced pluripotent stem cells (iPSC) were maintained in mTeSR1 with supplement. iPSCs were clump dissociated and passaged using enzyme-free Cell Dissociation Buffer. HEK293xT cells were maintained in DMEM, high glucose supplemented with 10% fetal bovine serum, heat inactivated. HEK293xT cells were dissociated and passaged using TrypLE Express Enzyme dissociation buffer. Small molecule neural precursor cells (NPC) were maintained in DMEM/F12, GlutaMAX supplement, containing N-2 Supplement (1:100 v/v), CTS B27 Supplement, XenoFree (1:50 v/v), L-ascorbic acid (150 μM), CHIR99021 (3 μM), and purmorphamine (0.5 μM). NPCs were dissociated and passaged using Accutase solution. iPS-MNs were maintained in DMEM/F12, GlutaMAX supplement, containing N-2 Supplement (1:100 v/v), CTS B27 Supplement, XenoFree (1:50 v/v), L-ascorbic acid (150 μM), recombinant human BDNF (2 ng/mL), recombinant human CNTF (2 ng/mL), and recombinant human GDNF (2 ng/mL). IPS-MNs were dissociated and passaged using Accumax solution. All cell types were cultured in humidified incubators at 37°C, 5% CO_2_.

### Coating plates for cell culture maintenance

For HEK293xT cells, 10 cm tissue culture plates were coated with 4 ml aqueous solution of 0.001% w/v poly-D-lysine (PDL) hydrobromide and incubated for 5 min. The PDL was then aspirated and the plates washed once with sterile water. For human induced pluripotent stem cells and small molecule neural precursor cells (NPC), 10 cm tissue culture plates were coated with 6 mL of Matrigel (diluted 1:60 v/v in DMEM/F12, GlutaMAX supplement) and incubated for 60 min. The Matrigel was then aspirated and the plates seeded with cells. For iPS-MNs, 10 cm tissue culture plates were coated with 4 ml aqueous solution containing 0.001% w/v PDL hydrobromide and 0.001% w/v poly-L-ornithine (PLO) hydrobromide and incubated overnight. The PDL/PLO was then aspirated and the plates washed once with sterile water. Next the plates were coated with 6 ml of mouse laminin (20 μg/ml in NPC media without CHIR99021 and purmorphamine) and incubated overnight. Finally, the laminin was aspirated and the plates were seeded with cells.

### Coating plates for screening and secondary assays

For HEK293xT cells, 384-well plates were coated by adding 10 μL of 0.001% w/v poly-D-lysine (PDL) hydrobromide to each well and incubating overnight. The PDL was then aspirated and the wells washed once with sterile water. For small molecule neural precursor cells (NPC), 384-well plates were coated by adding 10 μL of 0.001% w/v PDL to each well and incubating overnight. The PDL was then aspirated and the wells washed once with sterile water. Next, 15 μL of mouse laminin (20 μg/ml in NPC media without CHIR99021 and purmorphamine) was added to each well and incubated overnight. Finally, the laminin was aspirated and the plates were seeded with cells. For iPS-MNs, 384-well plates were coated by adding 10 μl of 0.001% w/v PDL + 0.001% w/v poly-L-ornithine (PLO) hydrobromide to each well and incubating overnight. The PDL/PLO was then aspirated and the wells washed once with sterile water. Next, 15 μl of mouse laminin (20 μg/ml in NPC media without CHIR99021 and purmorphamine) was added to each well and incubated overnight. Finally, the laminin was aspirated and the plates were seeded with cells. Liquid handling was performed with a Hamilton Microlab STAR fluid handler.

### Screen assay for identifying SG-modulating compounds

Using a Hamilton Microlab STAR liquid handler, HEK293xT cells were plated at 7500 cells per well of 384-well plates in 20 μl of HEK293xT media, while the CV-B small molecule neural precursor cells (NPC) were plated at 12000 cells per well of 384-well plates in 20 μl of NPC media without CHIR99021 or purmorphamine. Both cell types were then incubated overnight. After overnight incubation, cells were then pre-treated with screen compounds: Using a Labcyte ECHO 555 Liquid Handler, compounds were spotted into the wells in biological duplicate at a final concentration of 10 μM, and cells were incubated with compounds for 60 min. After compound pre-treatment, cells were stressed by using the Hamilton fluid handler to add NaAsO_2_ diluted in 20 μl of DMEM to each well, to a final concentration of 500 μM or 250 μM of NaAsO_2_ for HEK293xT cells or NPCs, respectively. Cells were incubated with NaAsO_2_ for 60 min. Finally, cells were fixed by adding 24% paraformaldehyde in PBS to each well to a final concentration of 4% and incubated for 45 min at room temperature. After fixing, the plates were washed three times with PBS. Nuclei were stained with 30 μl DAPI (1:5000 v/v in PBS) per well and incubated overnight at 4°C, after which the plates were washed once with PBS. Finally, to preserve the plates 30 μl of 50% v/v glycerol in PBS was added to each well. Cells were imaged in this glycerol solution.

### Robotic imaging of screen plates

Screen plates were imaged using a Thermo Scientific CRS CataLyst-5 Express robotic plate loader coupled with a GE Healthcare IN Cell Analyzer 1000 plate imager. Four fields at the center of each well were imaged with a 10× objective through 460 nm and 535 nm emission filters for DAPI and G3BP1-GFP, respectively. Images at 460 nm and 535 nm were taken with 250 ms and 750 ms exposures, respectively.

### Automated image segmentation and feature quantification

SG screen and secondary assay images were segmented and image features quantified using a custom CellProfiler pipeline (*71*). Briefly, nuclei were segmented and identified in the DAPI fluorescence channel images using a diameter cutoff of 9-80 pixels for HEK293xT cells, small molecule neural precursor cells (NPC), and iPS-MNs. Cell bodies were then extrapolated by overlaying the GFP fluorescence channel images and tracing radially outward from the nuclei to the limits of the cytoplasmic G3BP1-GFP signal. The cell bodies were used as masks to eliminate imaging artifacts outside of cell boundaries, such as background fluorescence or dead cells. After masking, punctate structures were enhanced by image processing for speckle-like features that were 10 pixels in diameter for HEK293xT cells or 7 pixels in diameter for NPCs and IPS-MNs, and these punctate structures were then annotated as features such as G3BP1-GFP SG or TDP-43 foci. Finally, the total image area which was enclosed in each of the identified features (nuclei, punctate structures, or intersections of features) was calculated and output to spreadsheets.

### Statistical analysis of screen data

SG-modulating compounds were defined as hits that significantly decrease or increase one or more of three metrics:

1. Amount of SG formation per cell, defined as the total image area enclosed in SGs divided by the total image area enclosed in nuclei (SG area/nuclei area).
2. Number of SGs per cell, defined as the number of SG puncta divided by the total image area enclosed in nuclei (SG count/nuclei area).
3. Average size of SGs, defined as the total image area enclosed in SGs divided by the number of SG puncta (SG area/SG count).

To identify hit compounds for these three metrics, a custom computational pipeline, Statistical Workflow for Identification of Molecular Modulators of ribonucleoprotEins by Random variance modeling (SWIMMER; open source and available publicly on yeolab github page), was built following standard statistical practices for high-throughput screening (*72, 73*). The pipeline begins by interpolating sporadic missing values via K-nearest neighbors missing data imputation, using K = 10 nearest neighbors and Euclidean distances (*74*). To reduce batch effects, the pipeline then performs a two-way median polish, alternating across rows and columns over 10 iterations (*72, 75*). Next, normalization is performed by computing b-scores, in which post-polish residuals are divided by the median absolute deviation of all post-polish residuals of the screen (*76*). To increase the power of hypothesis tests performed on the b-scores and therefore improve calling of screen hits, the screen was performed in biological duplicate, sample variances across the duplicate b-scores were computed, and an inverse gamma distribution was fitted to the distribution of sample variances (*77*). Fitting an inverse gamma distribution enables performing a modified one sample Student’s t-test which has increased statistical power for each screen compound (*72*). This modified t-test involves computing for each compound a modified t-statistic 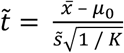, where 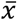 is the sample mean for the compound across *K* biological replicate b-scores, *μ*_0_ is the mean under the null hypothesis, and 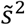 is the modified sample variance for the compound across *K* biological replicate b-scores. In the present SG screen, *μ*_0_ = 0, *K* = 2, and 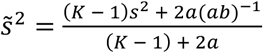, where *s*^2^ is the unmodified sample variance across *K* biological replicate b-scores and *a* and *b* are the fitted parameters of the inverse gamma distribution. Using these modified one sample Student’s t-tests, hits were called using a significance cutoff of α = 0.001; calling hits using a family-wise error rate proved too stringent and resulted in too many false negatives. Quality control (QC) for called hits was performed by examining the raw screen images for imaging artifacts or confounding effects such as autofluorescence or out-of-focus wells. Screen hits that passed QC were manually annotated with the compound names, skeletal formulae, and previously reported cellular targets of the compounds using the National Center for Biotechnology Information PubChem database.

### Computation of the screen assay Z-factor

To estimate whether the screen assay has adequate sensitivity and specificity for detecting hits in a high-throughput screening paradigm, the Z-factor was computed for the screen assay in each cell type: *Z* − *factor* = 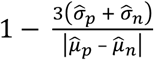, where 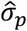 and 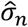 are the sample standard deviations for the positive and negative controls, respectively, and 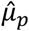 and 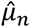 are the sample means for the positive and negative controls, respectively. The Z-factor is therefore a measure of the signal-to-noise in the screen assay, and an assay is suitable for high-throughput screen if the Z-factor is greater than 0.5 (*37*).

### Counterscreen assays

384-well plates were coated and cells were seeded as described above for 384-well screening plates. For dose response assays, hit compounds diluted in 20 μl DMEM were added to cells to final concentrations of 30, 10, 3.3, 1.1, 0.37, or 0.12 μM, and cells were incubated with compounds for 60 min. Cells were then stressed by adding NaAsO_2_ diluted in 40 μl of DMEM to each well, to a final concentration of 500 μM or 250 μM for HEK293xT cells or CV-B small molecule neural precursor cells (NPC), respectively. Cells were incubated with NaAsO_2_ for 60 min at 37°C, 5% CO_2_, after which cells were fixed, DAPI stained, and imaged. For heat shock stress assays, hit compounds diluted in 20 μl DMEM were added to cells to final concentrations of 10 μM, and cells were incubated with compounds for 60 min. Cells were then heat shocked by incubating for 60 min at 43°C, 5% CO_2_, after which cells were fixed, DAPI stained, and imaged. For thapsigargin stress assays, hit compounds diluted in 20 μL DMEM were added to cells to final concentrations of 10 μM, and cells were incubated with compounds for 60 min. Cells were then stressed by adding thapsigargin diluted in 40 μL DMEM to a final concentration of 50 μM or 1 μM for HEK293xT cells or NPCs, respectively. Cells were then incubated with thapsigargin for 60 min at 37°C, 5% CO2, after which cells were fixed, DAPI stained, and imaged. For the NaAsO_2_ pre-stress assays, cells were first stressed by adding NaAsO_2_ diluted in 20 μl DMEM to a final concentration of 500 μM or 250 μM for HEK293xT cells or NPCs, respectively. Cells were incubated with NaAsO_2_ for 60 min. Then, hit compounds diluted in 40 μl DMEM were added to cells to a final concentration of 10 μM, and cells were incubated with compounds for 60 min at 37°C, 5% CO_2_, after which cells were fixed, DAPI stained, and imaged.

### Estimating IC50s of SG inhibiting compounds

IC50s were estimated from logistic functions fitted to the dose response data of SG inhibiting compounds. Least squares regression was used to fit logistic functions of the type: 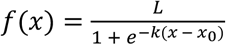, where *L* is the maximum value of the curve, *k* is the steepness of the curve and *x*_0_ is the midpoint of the curve. From these fitted logistic functions, the IC50 estimates are therefore the fitted x_0_ parameters, representing the midpoints of the dose-response transfer curves.

### Western blotting and quantification of eIF2α serine 51 phosphorylation

12-well plates were coated as described above for 384-well primary screening plates but scaled to 12-well plates. 2*10^5^ HEK293xT cells or CV-B small molecule neural precursor cells (NPC) were seeded into each well and incubated overnight. Screen hit compounds diluted in DMEM were added to cells to final concentrations of 10μM and incubated with cells for 60 min. Cells were then stressed by adding NaAsO_2_ diluted in DMEM to final concentrations of 500 μM or 250 μM for HEK293xT cells or NPCs, respectively, and incubated for 40 min. Next, the media was aspirated and cells were washed once with PBS at room temperature. Cells were lysed for 10 min on ice with 100 μL per well of ice-cold RIPA Buffer Solution (Teknova) supplemented with cOmplete, EDTA-free Protease Inhibitor Cocktail (1 tab/50 ml), PhosSTOP phosphatase inhibitor cocktail (1 tab/10 mL), and Benzonase nuclease cocktail (1:1000). A Pierce BCA Protein Assay Kit was used to quantify protein concentrations in lysates and 1 μg of total protein from each lysate was loaded for gel electrophoresis and Western blotting. Western blot membranes were blocked with 1% bovine serum albumin (BSA) in TBST for 60 min at room temperature, probed for serine 51 phosphorylate eIF2α using rabbit anti-EIF2S1 (Phospho-Ser51) primary antibody diluted 1:1000 in 0.5% BSA in TBST overnight at 4°C, and probed with secondary antibody diluted 1:2000 in 0.5% BSA in TBST for 60 min at room temperature. As control, total eIF2α was probed using rabbit anti-EIF2S1 primary antibody. Western blot band sizes were quantified by densitometry in ImageJ: Equally sized boxes were drawn around bands and pixel intensities were integrated over the box areas to give the band sizes (*78*).

### Isolation of SG-enriched fractions

SG-enriched fractions were isolated as described previously (*10*). Briefly, cells were grown in 10 cm plates to 75-90% confluency and stressed with NaAsO_2_ as described above. The media was then aspirated, cells were washed once with PBS at room temperature, and then lysed for 10 minutes on ice with 6 mL ice-cold NP-40 lysis buffer per 10 cm plate; formulation of this lysis buffer is as previously described (*10*). Lysed cells were scraped and collected in 15 ml centrifuge tubes in 3 ml aliquots. Lysates were sonicated using a Diagenode Bioruptor Plus for two cycles of 10 s on low power and one cycle of 10 s on high power. After each sonication cycle, lysates were chilled on ice for 20 s. Following sonication, the lysates were centrifuged at 1000 g for 5 min at 4°C to pellet nuclei. The supernatants were then collected into 1.5 ml centrifuge tubes and centrifuged at 18000 g for 20 min at 4°C to pellet SGs. The pellets were resuspended by trituration in 1/10 of the original volume of ice-cold NP-40 lysis buffer. The resuspended samples were finally centrifuged at 850 g for 2 min at 4°C to pellet membrane-bound organelles and other large debris. The remaining supernatants were collected as the SG-enriched fractions. The same fractionation procedure was performed on unstressed cells as a control.

### Western blotting and quantification of RBPs in SG-enriched fractions

A Pierce BCA Protein Assay Kit was used to quantify protein concentrations in SG-enriched fractions and 1 μg of total protein from each sample was loaded for gel electrophoresis and Western blotting. For input controls, 1 μg of total protein from whole cell lysates was also loaded. After transfer and blotting, Western blot band sizes were quantified by densitometry in ImageJ as described above. SG-enriched fraction band sizes were normalized by dividing by the band sizes of the corresponding input controls.

### Immunofluorescence probing and microscopy of SG-enriched fractions

In each well of a 96-well plate, 2 μg of total protein from SG-enriched fractions was added to 40 μl of ice-cold NP-40 lysis buffer and SGs in these fractions were allowed to settle by incubating overnight at 4°C. To fix the SGs, 8% paraformaldehyde (PFA) in PBS was added to each well to a final concentration of 4% PFA and plates were incubated for 10 min at room temperature. The PFA was aspirated and the wells were washed three times with PBS at room temperature. Next the wells were simultaneously permeabilized and blocked by adding 80 μL of 0.1% v/v Triton X-100 + 5% v/v serum (of the same species as secondary antibody) in PBS to each well and incubating the plates for 45 min at room temperature. The plates were then washed once with wash buffer (0.01% v/v Triton X-100 in PBS) before probing with primary antibody. Primary antibody was diluted in wash buffer containing 1% w/v bovine serum albumin (BSA). 40 μl of primary antibody solution was added per well and plates were overnight at 4°C. Plates were then washed five times with wash buffer. Secondary antibody was diluted in wash buffer containing 1% w/v BSA. 40 μl of secondary antibody solution was added per well and plates were incubated overnight at 4°C. Plates were washed 10 times with wash buffer and preserved by adding 80 μl of 50% v/v glycerol in PBS per well. Plates were then imaged at 20× magnification on a ZEISS Axio Vert.A1 inverted microscope.

### Quantifying co-localization of SGs and immunofluorescence-probed RBPs in SG-enriched fractions

Images were first pre-processed in ImageJ, to subtract background using the rolling ball method with a radius of 50 pixels (required for subsequent Manders Correlation Coefficient calculations), and then to transform the dynamic range of the brightness to 2-10 (arbitrary units). Co-localization of G3BP1-GFP SGs with RBPs probed with Alexa-555 conjugated antibodies was then quantified using Manders’ algorithm in the Fiji plugin Coloc2 to calculate the unthresholded Manders’ Correlation Coefficient between the GFP fluorescence channel image and the Alexa-555 fluorescence channel image (*79, 80*). In this way, co-localization is defined as the fraction of G3BP1-GFP-positive pixels that coincide with Alexa-555-positive pixels.

### Disrupting SG-RBP association in SG-enriched fractions

For digestion of SG-enriched fractions with RNase I, 10 μg of total protein from SG-enriched fractions was incubated with 0, 2, 8, or 32 Units of RNase I in 1.5 mL centrifuge tubes and shaken at 1200 rpm for 5 min at 37°C using an Eppendorf Thermomixer R. For incubation with 1,6-hexanediol, 10 μg of total protein from SG-enriched fractions was incubated with 1,6-hexanediol to a final concentration of 5% and nutated overnight at 4°C using a Fisherbrand Mini-Tube Rotator. For incubation with compounds, 10 μg of total protein from SG-enriched fractions was incubated with a screen hit compound to a final concentration of 100 μM and nutated overnight at 4°C using a Fisherbrand Mini-Tube Rotator. After overnight incubations, the SG-enriched fractions were centrifuged at 18000 × g for 20 min at 4°C to pellet remaining SG-like structures. The pellets were resuspended in equal volume lysis buffer and analyzed by Western blotting and immunofluorescence probing as described above.

### Disrupting TDP-43ΔNLS (86-414) association with SGs in H4 cells

H4 cells were stably transduced with lentivirus encoding G3BP1-mCherry and doxycycline-inducible GFP-TDP-43ΔNLS (86-414) made from pLVX-CMV_IRES-Hygro (Clontech) and pLVX-TetOne_IRES-Puro (Clontech), respectively. Cells were maintained in DMEM, high glucose supplemented with 10% fetal bovine serum, heat inactivated, and grown on 10 cm PDL coated plates, prepared as described above. H4 cells were dissociated and passaged using TrypLE Express Enzyme dissociation buffer. 2*10^5^ cells in 50 μL media were seeded into each well of a 96-well plate (Perkin-Elmer). Following overnight incubation in a humidified incubator at 37°C, 5% CO_2_, the cells were induced with doxycycline (1 ng/mL) for 24 hours. After doxycycline induction, hit compounds diluted in 50 μl media were added to final concentrations of 5 μM, and cells were incubated with compounds for 30 min. Cells were then stressed by adding NaAsO_2_ or thapsigargin diluted in 100 μl media to a final concentration of 500 μM or 5 μM, respectively, and incubated for 60 min. Cells were then fixed, stained with DAPI, and imaged at 40× magnification on an Opera Phenix high content imaging system (Perkin Elmer).

### IPS-MN puromycin stress and recovery assays

iPS-MNs were generated from human induced pluripotent stem cells as described above. Day 28 iPS-MNs were dissociated into single cells by adding 3 mL Accumax into each well of a 6-well plate of IPS-MNs and incubating for 60 min. The iPS-MNs were then gently triturated 10-15 times with a P1000 pipette, and then incubated for a further 15 min at 37°C, 5% CO_2_ to complete the dissociation. The IPS-MNs were transferred to 15 ml centrifuge tubes containing 6 ml DMEM/F12, GlutaMAX supplement and cells were pelleted by centrifuging for 4 min at 200× g. The cell pellet was then resuspended in 1 ml IPS-MN media supplemented with 10 μM Y-27632 (media formulation described above) by gently triturating 10-20 times with a P1000 pipette, and cell clumps were filtered out using a 40 μm filter. 10^4^ iPS-MNs were seeded into each well of a laminin-coated 384-well plate; plate coating was as described above. Cells were allowed to adhere by incubating overnight at 37°C, 5% CO_2_ and then stressed by adding puromycin diluted in 20 μl of IPS-MN media to a final concentration of 5 μg/ml. After incubation with puromycin for 24 hours, three half-media changes with IPS-MN media were performed to wash out the stressor. The MNs were then incubated for 24 hours at 37°C, 5% CO_2_, after which cells were fixed and DAPI stained. IPS-MNs were also fixed at intermediate time points to generate a time course for SG and TDP-43 foci assembly/disassembly dynamics during puromycin stress and stress recovery. G3BP1 and TDP-43 were probed by immunofluorescence according to the procedure described above for immunofluorescence probing of SG-enriched fractions. IPS-MNs were imaged as described above for imaging 384-well primary screening plates. SG-enriched fractions were also isolated from IPS-MNs following the protocol described above.

### Hit compound inhibition of NaAsO_2_, thapsigargin, or puromycin-induced SGs in iPS-MNs

IPS-MNs were differentiated and seeded into pre-coated 384-well plates as described above. IPS-MNs were allowed to adhere by incubating for 48 hours at 37°C, 5% CO_2_; IPS-MNs were fed 24 hours after seeding by adding 20 μL of IPS-MN media to each well to a final volume of 40 μL per well. For NaAsO_2_ stress assays, hit compounds diluted in 20 μL media were added to IPS-MNs to a final concentration of 10 μM, and MNs were incubated with compounds for 80 min. IPS-MNs were then stressed by adding NaAsO_2_ diluted in 20 μL IPS-MN media to a final concentration of 100 μM and incubated for 120 min at 37°C, 5% CO_2_, after which cells were fixed. For thapsigargin stress assays, hit compounds diluted in 20 μl MN media were added to MNs to final concentrations of 10 μM, and MNs were incubated with compounds for 80 min. MNs were then stressed by adding thapsigargin diluted in 20 μl MN media to a final concentration of 250 nM and incubated for 120 min at 37°C, 5% CO_2_, after which cells were fixed. For puromycin stress assays, hit compounds and puromycin were diluted in 40 μl MN media and added to MNs to final compound concentrations of 5 μM and final puromycin concentration of 5 μg/ml. MNs were incubated with compounds and puromycin for 12 hours at 37°C, 5% CO_2_, after which cells were fixed. G3BP1 and TDP-43 were probed by immunofluorescence according to the procedure described above for immunofluorescence probing of SG-enriched fractions and nuclei were stained with DAPI. MNs were imaged as described above for imaging 384-well primary screening plates.

| REAGENT or RESOURCE | SOURCE | IDENTIFIER |
| --- | --- | --- |
| **Antibodies** |  |  |
| Rabbit anti-ATAXIN2 | Proteintech | 21776-1-AP |
| Rabbit anti-CAPRIN1 | Proteintech | 15112-1-AP |
| Mouse anti-CYC | Santa Cruz Biotechnology | sc-13560 |
| Rabbit anti-DDX3 | Bethyl | A300-474A |
| Rabbit anti-DDX6 | Abcam | ab40684 |
| Mouse anti-EIF2S1 | Santa Cruz Biotechnology | sc-133132 |
| Rabbit anti-EIF2S1 (Phospho-Ser51) | Abcam | ab32157 |
| Rabbit anti-EIF4G1 | MBL | RN002P |
| Rabbit anti-FUS/TLS | Abcam | ab70381 |
| Rabbit anti-G3BP1 | MBL | RN048PW |
| Mouse anti-GFAP | Millipore Sigma | NE1015 |
| Mouse anti-HB9 | Santa Cruz Biotechnology | sc-515769 |
| Mouse anti-HNRNPA1 | Novus | NB100-672 |
| Rabbit anti-HNRNPA2B1 | GeneTex | GTX127928 |
| Mouse anti-ISL1 | Abcam | ab86501 |
| Rabbit anti-NANOG | Abcam | ab80892 |
| Mouse anti-O4 | Millipore Sigma | MAB345 |
| Rabbit anti-OLIG2 | Millipore Sigma | AB9610 |
| Rabbit anti-PABPC1 | Abcam | ab21060 |
| Mouse anti-SMI-31 | Millipore Sigma | NE1022 |
| Mouse anti-TARDBP | Abnova | H00023435-M01 |
| Goat anti-TIA1 | Santa Cruz Biotechnology | SC-1751 |
| Chicken anti-TUBB3 | Abcam | ab107216 |
| Rabbit anti-UBAP2L | Bethyl | A300-533A-M |
| Rabbit anti-USP10 | Abcam | ab109219 |
| Goat anti-Rabbit IgG, HRP conjugate | ThermoFisher Scientific | 31460 |
| Goat anti-Mouse IgG, HRP conjugate | Millipore Sigma | 401215 |
| Rabbit anti-Goat IgG, HRP conjugate | Millipore Sigma | 401515 |
| Goat anti-Rabbit IgG, Alexa Fluor 555 | ThermoFisher Scientific | A21429 |
| Goat anti-Mouse IgG, Alexa Fluor 555 | ThermoFisher Scientific | A21422 |
| Donkey anti-Goat IgG, Alexa Fluor 555 | ThermoFisher Scientific | A21432 |
| **Chemicals, Peptides, and Recombinant Proteins** |  |  |
| Sodium (meta)arsenite | Millipore Sigma | 1062771000 |
| Thapsigargin | Tocris Bioscience Bioscience | 1138 |
| DMSO | Millipore Sigma | D2650 |
| UCSF Neuroprotective Library | UCSF | N/A |
| Genentech Neuroprotective Library | Genentech | N/A |
| US Drug Collection | MicroSource Discovery Systems | N/A |
| International Drug Collection | MicroSource Discovery Systems | N/A |
| Spectrum Collection | MicroSource Discovery Systems | N/A |
| LOPAC1280 | Millipore Sigma | N/A |
| Prestwick Chemical Library | Prestwick | N/A |
| 8-hydroxyquinoline | Santa Cruz Biotechnology | sc-202896 |
| Anacardic acid | Santa Cruz Biotechnology | sc-202463A |
| Anisomycin | Millipore Sigma | A9789 |
| Benzethonium | Millipore Sigma | B8879 |
| CA43/WS3 | Millipore Sigma | SML0758 |
| Cycloheximide | Tocris Bioscience Bioscience | 970 |
| Daunorubicin | Santa Cruz Biotechnology | sc-200921 |
| Digitoxin | Santa Cruz Biotechnology | sc-207577A |
| Digoxin | Millipore Sigma | D6003 |
| Doxorubicin | Fisher Scientific | ICN15910110 |
| Gitoxigenin | Millipore Sigma | S461954 |
| GSK2606414 (PERK inhibitor) | Apexbio | A3448 |
| ISRIB | Millipore Sigma | SML0843 |
| Kenpaullone | Tocris Bioscience Bioscience | 1398 |
| Mitoxantrone | Santa Cruz Biotechnology | sc-203136A |
| Ouabain | Millipore Sigma | O3125 |
| Paclitaxel | Santa Cruz Biotechnology | sc-201439 |
| Pararosaniline pamoate | Millipore Sigma | P3750 |
| PD98059 (ERK inhibitor) | Santa Cruz Biotechnology | sc-3532 |
| Penfluridol | Santa Cruz Biotechnology | sc-255410 |
| Proscillaridin A | Santa Cruz Biotechnology | sc-202775 |
| Pyrvinium pamoate | Millipore Sigma | P0027 |
| Quinacrine | Santa Cruz Biotechnology | sc-204222 |
| ST1 | UCSF | N/A |
| Tyrphostin A9 | Santa Cruz Biotechnology | sc-200568 |
| mTeSR1 with supplement | Stemcell Technologies | 85850 |
| SB431542 | Tocris Bioscience | 1614 |
| Dorsomorphin | Tocris Bioscience | 3093 |
| DMEM, high glucose | Gibco | 11965092 |
| Fetal Bovine Serum, qualified, heat inactivated, US origin | Gibco | 16140071 |
| MEM Non-Essential Amino Acids Solution (100X) | Gibco | 11140050 |
| L-Glutamine (200 mM) | Gibco | A2916801 |
| DMEM/F12, GlutaMAX supplement | Gibco | 10565018 |
| N-2 Supplement | Gibco | 17502048 |
| CTS B27 Supplement, XenoFree | Gibco | A1486701 |
| L-Ascorbic acid | Millipore Sigma | A4544 |
| CHIR99021 | Tocris Bioscience | 4423 |
| Purmorphamine | Tocris Bioscience | 4551 |
| Y-27632 | Tocris Bioscience | 1254 |
| Smoothened Agonist, SAG | Millipore Sigma | 566660 |
| Retinoic acid | Millipore Sigma | R2625 |
| DAPT | Tocris Bioscience | 2634 |
| Recombinant human BDNF | PeproTech | 450-02 |
| Recombinant human CNTF | PeproTech | 450-13 |
| Recombinant human GDNF | PeproTech | 450-10 |
| Cell Dissociation Buffer, enzyme-free, PBS | ThermoFisher Scientific | 13151014 |
| TrypLE Express Enzyme (1X), no phenol red | ThermoFisher Scientific | 12604013 |
| Accutase | Innovative Cell Technologies | AT104 |
| Accumax | Innovative Cell Technologies | AM105 |
| Matrigel, hESC-Qualified | Corning | 354277 |
| Laminin Mouse Protein, Natural | ThermoFisher Scientific | 23017015 |
| Poly-D-lysine hydrobromide | Millipore Sigma | P6407 |
| Poly-L-ornithine hydrobromide | Millipore Sigma | P3655 |
| Paraformaldehyde | Millipore Sigma | P6148 |
| Triton X-100 | Millipore Sigma | X100 |
| Bovine Serum Albumin | Millipore Sigma | A8806 |
| Goat Serum | Millipore Sigma | G9023 |
| Donkey Serum | Millipore Sigma | D9663 |
| DAPI | Millipore Sigma | D9542 |
| Glycerol | Millipore Sigma | G5516 |
| Antifoam B | Millipore Sigma | A5757 |
| Heparin | Millipore Sigma | H3149 |
| IGEPAL CA-630 | Millipore Sigma | I8896 |
| RIPA Buffer Solution | Teknova | R3792 |
| Benzonase | Millipore Sigma | E1014 |
| RNase I | ThermoFisher Scientific | AM2295 |
| Tween 20 | Millipore Sigma | P1379 |
| 96-well optical bottom microplate | ThermoFisher Scientific | 165305 |
| 384-wl cell culture microplate | Greiner Bio-One | 781091 |
| NuPAGE LDS Sample Buffer | ThermoFisher Scientific | NP0007 |
| **Experimental Models: Cell Lines** |  |  |
| HEK293xT | Clontech | 632180 |
| HEK293xT G3BP1-GFP | This paper | N/A |
| CV-B iPSC | (Gore et al., 2011) | N/A |
| CV-B iPSC G3BP1-GFP | This paper | N/A |
| CV-B smNPC G3BP1-GFP | This paper | N/A |
| CV-B MN G3BP1-GFP | This paper | N/A |
| Kin1ALS6.3 Control iPSC | (Krach et al., 2018) | N/A |
| Kin1ALS6.2 Control iPSC | (Krach et al., 2018) | N/A |
| Kin2ALS6.4 Control iPSC | (Krach et al., 2018) | N/A |
| Kin1ALS17.3 Control iPSC | (Krach et al., 2018) | N/A |
| Kin3ALS6.4 TDP-43 N352S Mutant iPSC | (Krach et al., 2018) | N/A |
| Kin2ALS17.5 TDP-43 N352S Mutant iPSC | (Krach et al., 2018) | N/A |
| ALS17.5 TDP-43 N352S Mutant iPSC | (Krach et al., 2018) | N/A |
| 47d TDP-43 G298S Mutant iPSC | (Alami et al., 2014) | N/A |
| GY6.2 FUS R521G Mutant iPSC | (Kapeli et al., 2016) | N/A |
| GY7.3 FUS R521G Mutant iPSC | (Kapeli et al., 2016) | N/A |
| Kin1ALS6.3 Control MN | This paper | N/A |
| Kin1ALS6.2 Control MN | This paper | N/A |
| Kin2ALS6.4 Control MN | This paper | N/A |
| Kin1ALS17.3 Control MN | This paper | N/A |
| Kin3ALS6.4 TDP-43 N352S Mutant MN | This paper | N/A |
| Kin2ALS17.5 TDP-43 N352S Mutant MN | This paper | N/A |
| ALS17.5 TDP-43 N352S Mutant MN | This paper | N/A |
| 47d TDP-43 G298S Mutant MN | This paper | N/A |
| GY6.2 FUS R521G Mutant MN | This paper | N/A |
| GY7.3 FUS R521G Mutant MN | This paper | N/A |
| **Critical Commercial Assays** |  |  |
| CytoTune-iPS Sendai Reprogramming Kit | ThermoFisher Scientific | A1378001 |
| PhosSTOP | Millipore Sigma | 4906837001 |
| cComplete, EDTA-free Protease Inhibitor Cocktail | Millipore Sigma | 5056489001 |
| Pierce BCA Protein Assay Kit | ThermoFisher Scientific | 23225 |
| NuPAGE 4-12% Bis-Tris Midi Protein Gels | ThermoFisher Scientific | WG1403BOX |
| **Recombinant DNA** |  |  |
| GFP targeting vector | System Biosciences | HR120PA-1 |
| pSpCas9(BB)-2A-GFP (PX458) | Gift from Feng Zhang | Addgene plasmid # 48138 |
| pLVX-CMV_IRES-Hygro | Clontech | 632182 |
| pLVX-TetOne_IRES-Puro | Clontech | 631847 |
| **Software and Algorithms** |  |  |
| Image segmentation with CellProfiler | (Carpenter et al., 2006) | http://www.cellprofiler.org/ |
| Identifying hits with random variance modeling | This paper | N/A |
| Image colocalization with ImageJ | (Schindelin et al., 2012) | https://imagej.net/Coloc_2 |

